# Rapid evolution of synteny associated with multiple origins of dioecy and XY sex determination

**DOI:** 10.64898/2025.12.21.695719

**Authors:** Mark S. Hibbins, Cassandre Pyne, Mykhailo Sukmaniuk, Emily Glasgow, Meng Yuan, Yunchen Gong, Spencer C.H. Barrett, Stephen I. Wright

## Abstract

Chromosomal rearrangements are a major driver of evolutionary innovation, shaping processes including local adaptation, speciation, and sex chromosome evolution. Multispecies synteny datasets are rich in information on the drivers of genomic rearrangement, but statistical approaches that enable insights to be obtained from this information are still in their infancy. Here, we present a novel framework for the application of phylogenetic comparative methods to multispecies synteny datasets. We apply this approach to *Rumex*, a clade of flowering plants that exhibits rapid karyotypic evolution, including multiple origins of XY sex determination from hermaphroditic ancestors. Leveraging new genome assemblies, we find evidence for accelerated syntenic evolution associated with evolutionary transitions to dioecy, highlighting how explicit phylogenetic hypothesis testing can generate new insights into adaptive hypotheses for rearrangements.

---

Differences in karyotype are among the oldest recognized forms of genetic variation. Such variation is driven by mutations that change the structure, composition, and/or number of chromosomes. These rearrangements can affect a significant amount of the genome and have large functional and evolutionary consequences. Classic models describe scenarios under which rearrangements can be favored in a population, such as capturing beneficial haplotypes during local adaptation (*1-2*), extending the sex-linked regions of sex chromosomes (*3-5*), or reinforcing species barriers (*6-8*). While rigorous statistical frameworks modelling the evolution of nucleotide substitutions on phylogenies have existed for decades, analogous approaches have yet to be developed for larger-scale mutations such as rearrangements. For this reason, in addition to historical challenges in assembling whole-chromosome scaffolds in non-model species, empirical investigations of the role of structural changes in macroevolutionary processes have been remarkably limited.

Since the genomic age, the primary approach for detecting chromosomal rearrangements has been through the inference of synteny: the conservation of the physical ordering of genes along segments of chromosomes. Chromosomal rearrangements cause shifts in the physical adjacency of syntenic regions on a chromosome, enabling for reconstruction of ancestral karyotypes across species (*9-14*). Advances in accurate long-read sequencing technologies, in addition to new computational tools (*15*), enable the inference of syntenic regions in large multispecies datasets with high resolution. While rich in information, interpretation of these datasets has remained largely qualitative. Ancestral genome reconstruction, while powerful, is based on parsimony, which assumes minimal convergence and does not allow for statistically explicit hypothesis testing. Probabilistic approaches in an explicitly phylogenetic framework are needed to conduct rigorous hypothesis testing of the drivers of genomic rearrangement across species. For example, one may be interested in testing if phenotypic characters across a phylogeny are associated with broad shifts in the rate of genomic rearrangement, or accelerated rearrangement of specific chromosomes or syntenic regions.

In the flowering plant genus *Rumex* (Polygonaceae), chromosomal rearrangements are thought to have played a pivotal evolutionary role in species divergence. Dioecy has evolved multiple times in the genus, accompanied by successive reductions in chromosome number and at least two independent origins of XY sex determination in less than 20 MYA of divergence (*16-18*). Past work in dioecious *Rumex hastatulus* has revealed rapid rearrangement of both sex chromosomes and autosomes, including an X-autosome fusion leading to a within-species XY/XYY polymorphism, reciprocal Y translocations, and segregating inversions (*18-21*). However, apart from chromosome number variation, the history of rearrangement across the rest of the genus remains uncharacterized. Given the significant observed diversity of sexual systems, phylogenomic analyses of *Rumex* syntenic regions would provide new insights into classic theories on the role of chromosomal rearrangements in shaping sexual antagonism and sex determination. In particular, if chromosomal rearrangements spread adaptively on sex chromosomes in association with the resolution of sexual antagonism, we would predict that dioecious taxa should experience elevated rates of chromosomal rearrangements, particularly on the sex chromosomes.

In this study, we develop an approach based on the physical adjacency of syntenic regions that allows for phylogenetic comparative methods to be applied to multispecies synteny datasets to estimate rates of genomic rearrangement. We also implement a forward-time phylogenetic simulator of chromosomal rearrangements to help validate our approach. The method and simulator are incorporated in the new open-source R package *SynPhyMe* (<u>Syn</u>teny <u>Phy</u>logenetic <u>Me</u>thod). We generated ten new high-quality genome assemblies for six species of *Rumex*, and used *SynPhyMe* to uncover accelerated genome evolution associated with multiple transitions to dioecy from hermaphroditic ancestors in the genus. We also uncovered specific syntenic regions associated with transitions to dioecy and discuss their potential functional significance. Overall, our work provides new avenues for investigating the macroevolutionary drivers of genome diversification.

## Estimating rates of synteny evolution across phylogenies

*SynPhyMe* maps syntenic adjacencies to binary presence/absence phylogenetic characters for downstream inference with phylogenetic comparative methods. The input dataset is formatted based on the riparian plotting functions of GENESPACE (*15*), which phase syntenic regions to a reference genome and allow for tracking of custom genomic segments in the reference across species. The output of these functions can be given directly to *SynPhyMe*, along with an ultrametric phylogenetic tree of the relevant species, or a similar input file can be manually constructed from the output of other synteny inference software. As synteny inference is still conducted in a pairwise fashion between genomes, we assign adjacencies to each species based on the comparison with a nearest phylogenetic neighbor (Figure 1). The resulting output is an *n* by *m* binary character matrix, where *n* is the number of species and *m* is the number of adjacencies observed in at least one species (Figure 1).

**Figure 1:**
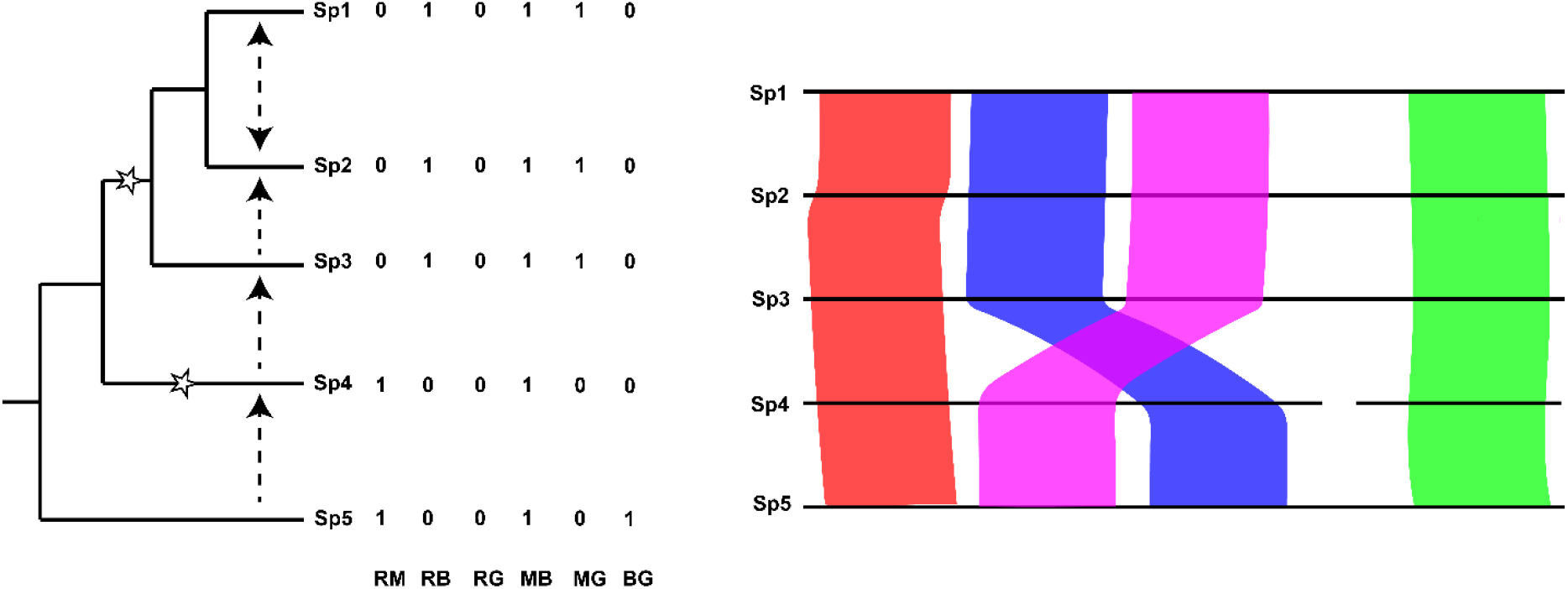
Mapping syntenic regions to phylogenetic characters. For a hypothetical phylogeny of five species, a hypothetical chromosome is divided into four segments in the reference genome (Sp5) and these segments are phased across the other species, giving six possible adjacencies. Two rearrangements occur along this phylogeny (labelled by stars on the relevant branches): a fission in Sp4 that separates the green region, and an inversion in the ancestor of Sp1, Sp2, and Sp3 that reverses the order of the magenta and blue regions. The comparison used to assign adjacencies to each species is indicated by the dashed arrows on the phylogeny.

For inference, we use the simplest instantiation of the Mk model (*22-23*) where a single rate parameter, *q*, models forward and backward character transitions. We use a composite likelihood approach where partial likelihoods are calculated for individual adjacencies using the pruning algorithm (*24*), and the log-sum of these likelihoods is optimized to obtain a genome-wide estimate of *q*. To validate this approach, we conducted standard phylogenetic simulations of binary characters under the Mk model, in addition to implementing in *SynPhyMe* a forward-time simulator of karyotype evolution driven by chromosomal rearrangements. Both simulation approaches highlight the accuracy of the method, with simulated value of *q* and the chromosomal mutation rate both being correlated with estimated values of *q* (Figure 2).

**Figure 2:**
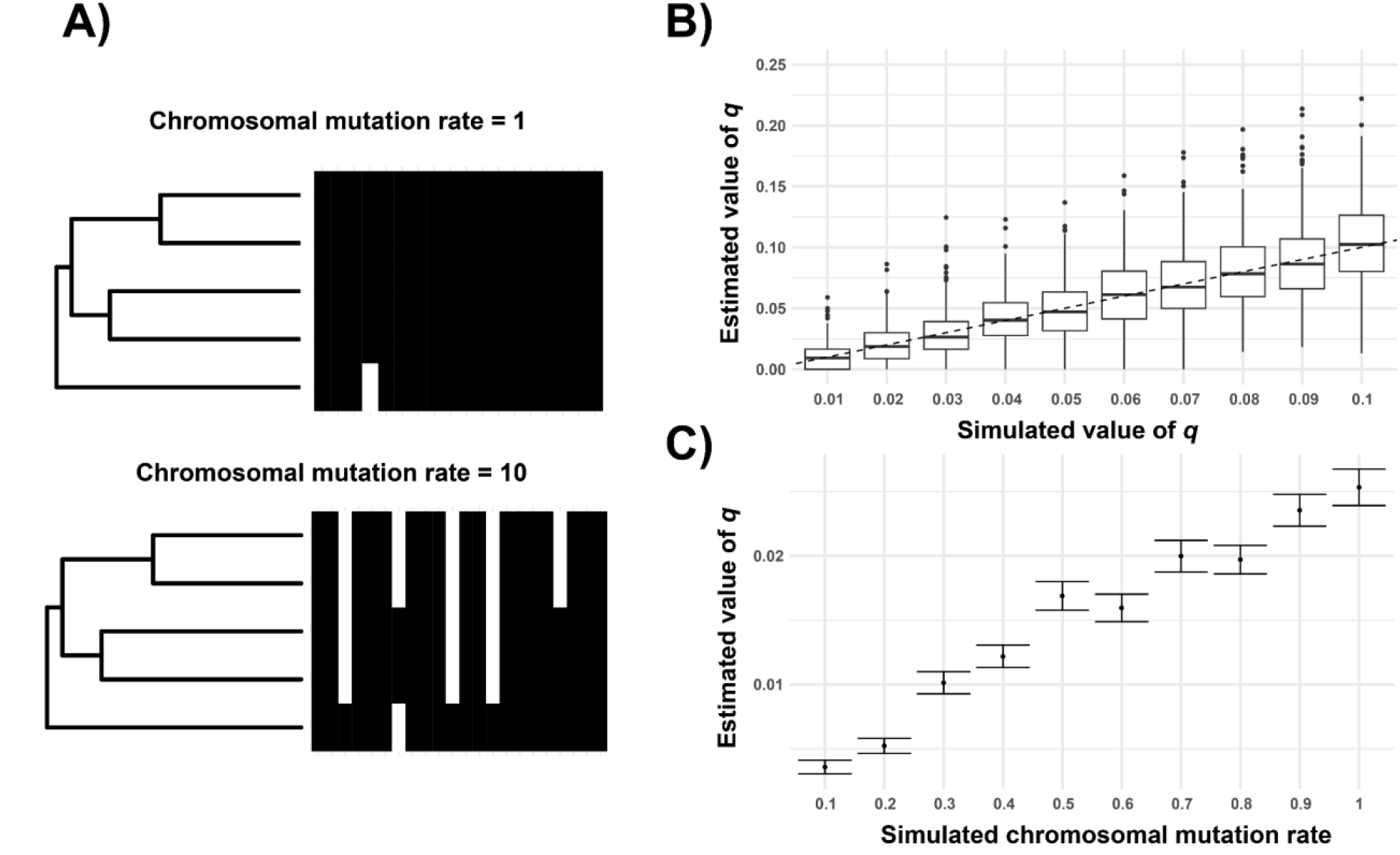
Evaluating our rate estimation approach with simulations. A) Example phylogenetic binary presence (black) / absence (white) matrices for syntenic adjacencies generated from our chromosomal rearrangement simulator. A representative output for a low mutation rate (top) and high mutation rate (bottom) are shown. B) Accuracy of our composite likelihood estimation of *q* on binary characters simulated from phylogenies under the MK model. Dashed line indicates the 1:1 relation between simulated and estimated values that would indicate perfect accuracy. C) Relationship between the per-branch length chromosomal mutation rate given to our rearrangement simulator, and the value of Q estimated from those simulations using our composite likelihood approach. Error bars indicate the standard error.

To enable evolutionary hypothesis testing, we also implemented a two-rate model where the user specifies a set of foreground branches on the phylogeny, and a rate multiplier *r* is estimated for these branches relative to the background *q* value of the rest of the tree. We use a simulation approach to evaluate the significance of this two-rate model against its nested one-rate model using the AIC criterion. This approach may be useful for users who wish to investigate if variation in the rate of syntenic rearrangement is associated with particular phenotypes. *SynPhyMe* can also return the likelihoods and parameters estimated for each individual adjacency, allowing users to identify the fastest evolving syntenic regions and investigate their functional significance. Finally, we note that beyond the analyses implemented here, users may also save the adjacency character matrix and use methods implemented in other packages that can handle multivariate phenotypic datasets (*25*).

## Rearrangement and sexual system shifts in *Rumex*

We generated long-read de novo genome assemblies and annotations for six *Rumex* species (Supplementary Figures 1-13, Supplementary Tables 1-2) and analyzed them alongside previously published assemblies for three species (*18, 21, 26*). We used GENESPACE to infer syntenic regions and estimated a new time-calibrated phylogeny using syntenic orthologs to reconstruct the history of rearrangements across the genus (Figure 3). Our phylogeny recapitulates previously inferred relationships, including two separate origins of dioecy, here denoted as the XYY and XY clades (*16, 18*). We uncovered a highly dynamic history of chromosomal rearrangement across the genus within a relatively short evolutionary timespan. Fissions, fusions, and translocations all occur frequently, affecting both autosomes and sex chromosomes in both hermaphroditic and dioecious species. Although the level of conservation varies, not a single chromosome in *R. salicifolius* is conserved across the whole phylogeny. Significantly, rearrangements occur in every syntenic comparison, even between the two *R. hastatulus* populations which are diverged by less than 1 MYA.

**Figure 3:**
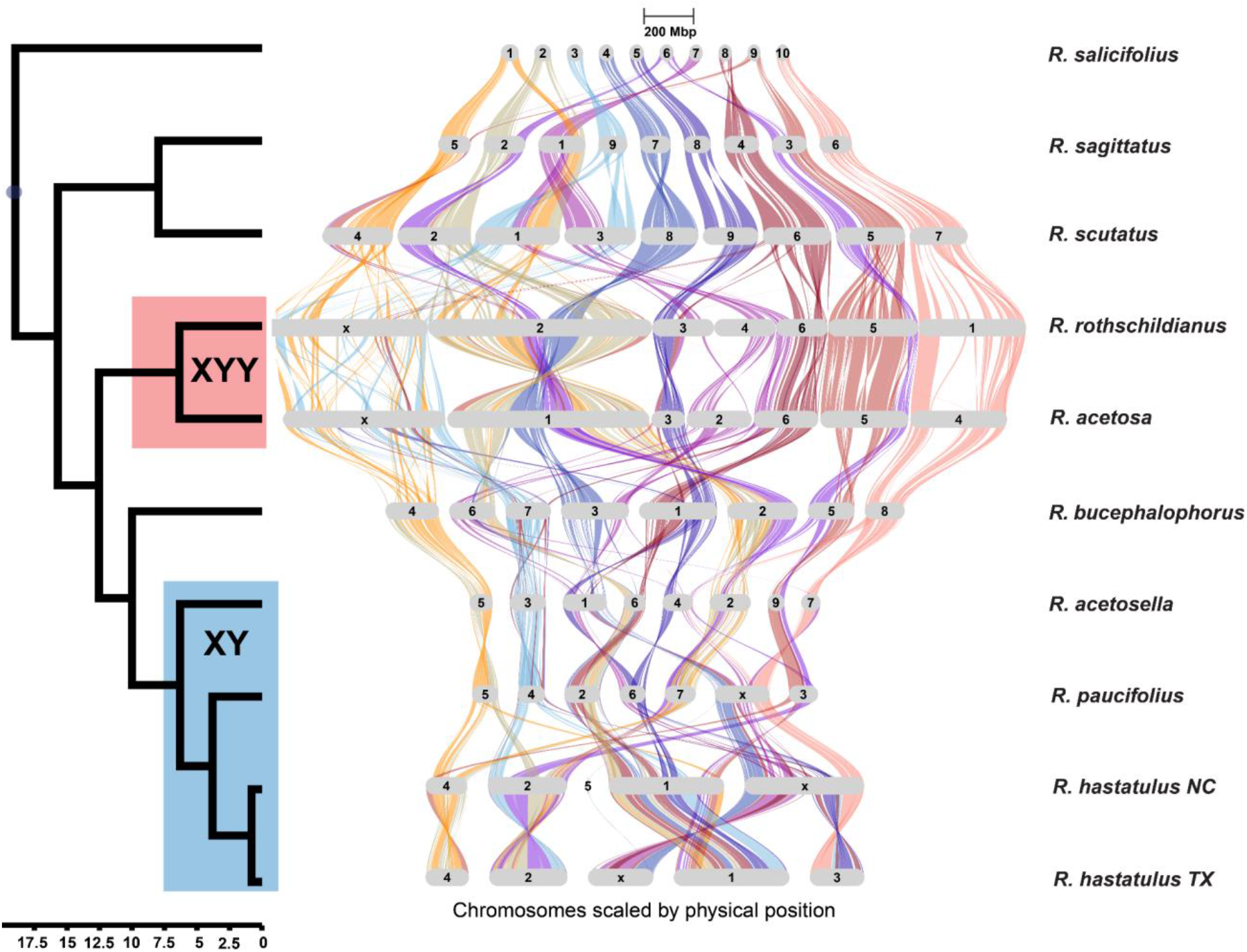
The evolution of synteny in *Rumex*. Riparian plot from GENESPACE tracking syntenic regions, aligned with our updated phylogeny inferred from syntenic orthologs. Color assignments in the riparian plot are based on synteny to individual chromosomes in the outgroup species, *R. salicifolius*. Chromosomes are scaled to physical size (scale bar at the top indicates size of 200 Mbp of sequence). Timescale under the phylogeny is in millions of years ago (MYA).

In the XYY clade (Fig. 3, red), our results are consistent with a single origin of the XYY system, though synteny has eroded significantly between the X chromosomes of *R. rothschildianus* and *R. acetosa* since their divergence ∼ 7 MYA (Figure 3). Consistent with theories of XYY system formation, the X appears to be primarily the product of a chromosomal fusion/translocation between regions syntenic to chromosomes 1 and 3 in *R. salicifolius*, in addition to containing a small shared segment syntenic to *R. salicifolius* chromosome 8, suggesting an additional smaller-scale insertion. These syntenic regions are interspersed throughout the X chromosomes, indicating numerous additional intra-chromosomal rearrangements following the initial fusion. Aside from the X, the chromosomes of the two XYY species are largely collinear. However, their genomes are significantly rearranged compared to the hermaphroditic species *R. scutatus*, potentially indicating an elevated rate of rearrangement in the ancestor of the two dioecious clades.

In the XY clade (Fig. 3, blue), we uncover a remarkable history of repeated rearrangements involving various combinations of sex-linked and autosomal regions, in less than 7 MYA of divergence. A shared sex-linked region, itself the product of multiple more ancestral autosomal rearrangements, appears to be present in all species (chromosome 1 is presumed to be the X in *R. acetosella*, but not confirmed). This region shares no synteny with the X chromosome of the XYY clade, confirming an independent origin of this system. In contrast to the canonical fusion model resulting in the XYY cytotype of eastern *R. hastatulus*, we find that an X-autosome fusion was likely ancestral to *R. paucifolius* and *R. hastatulus*, with an additional fission of the X producing autosome 3 of western *R. hastatulus*. It is unclear what led to the formation of the eastern XYY system, as it is not present in *R. paucifolius* despite sharing the same X-autosome fusion, but it may have involved an additional rearrangement incorporating part of *R. paucifolius* autosome 6. More explicit phylogenetic models of rearrangement processes will be necessary in the future to resolve the exact history of rearrangement in this group with confidence.

## Convergence and the genetic architecture of sex-linked regions

Theory predicts elevated rates of rearrangement involving sex chromosomes (*3, 27*) because rearrangements can capture haplotypes containing sexually antagonistic variation and link them to sex-linked regions. We used the two-rate method implemented in *SynPhyMe* to test if dioecious *Rumex* species (highlighted in Fig. 3) rearrange their genomes faster than their hermaphroditic relatives. We found that the two-rate model was a significantly (*p* < 0.01) better fit to the data, with parameter estimates of *q* = 0.00466 and *r* = 11.8, implying a nearly 12x faster rate of rearrangement in the dioecious lineages (Figure 4). We then used phylogenetic regression to identify syntenic regions repeatedly involved in rearrangement in dioecious lineages, identifying three marginally significant (p < 0.1) adjacencies all containing segments of *R. salicifolius* chromosome 8 (Supplementary Table 3). This chromosome is the only ancestral chromosome in *R. salicifolius* with regions syntenic to all X chromosomes in our dataset (Supplementary Figure 1), suggesting a potentially important functional role for the establishment of sex-linked regions.

**Figure 4:**
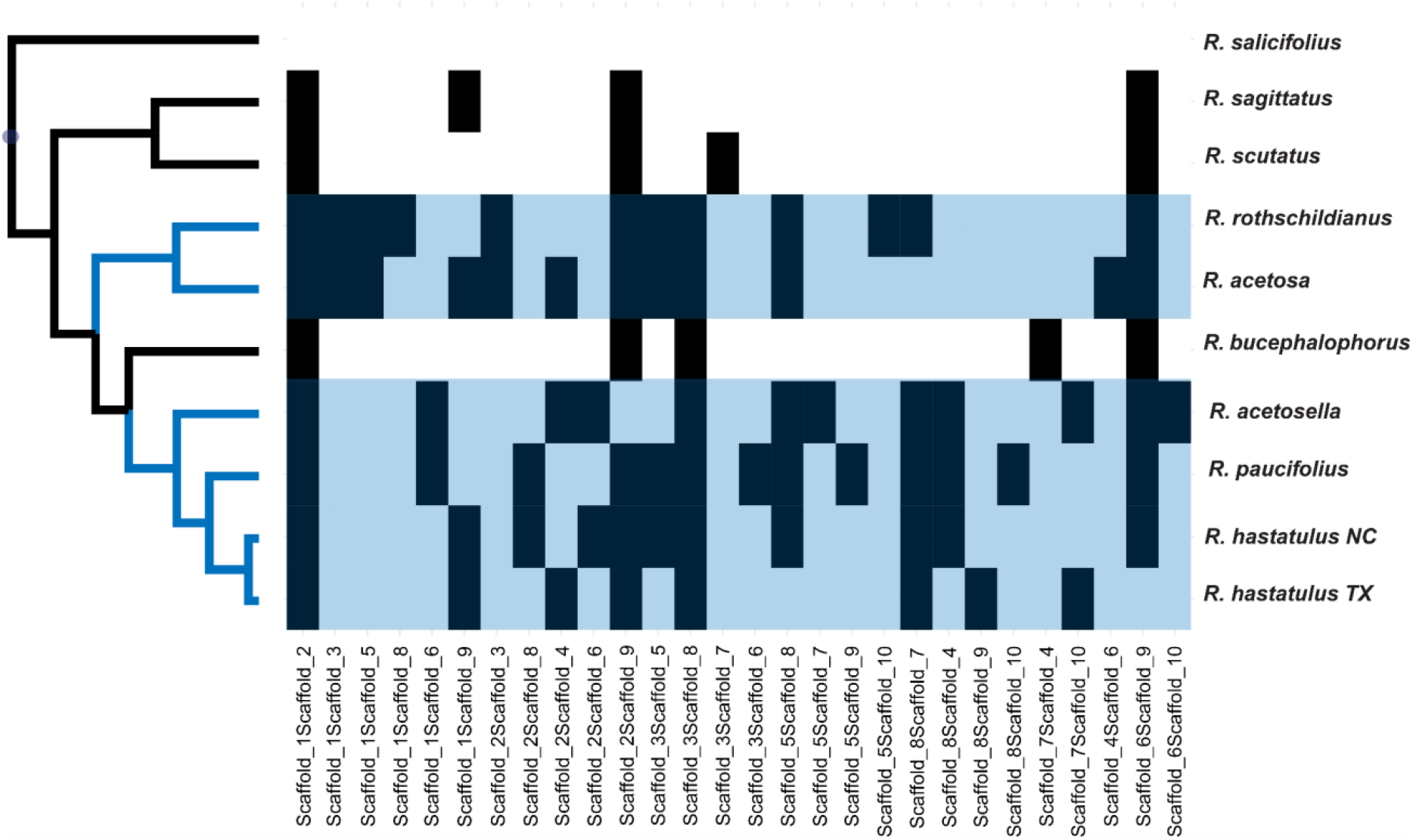
Rapid evolution of syntenic adjacencies in dioecious *Rumex* species. Binary presence-absence matrix (black = present) of observed syntenic adjacencies (columns) across species in our time-calibrated phylogeny of *Rumex* (rows). No adjacencies are present in *R. salicifolius* as they are defined with respect to individual chromosomes in this species. Phylogenetic branches, and their associated syntenic adjacencies, where we estimated an 11.8x faster rate of syntenic evolution relative to the rest of the tree are labelled in blue.

While some form of this syntenic region was present on all X chromosomes, individual genes within these regions do not appear to be repeatedly recruited or retained. We found no significant enrichment (*p* = 0.1702) of *R. salicifolius* chromosome 8 genes on the X chromosomes. Only two genes were found on two X chromosome regions syntenic to *R. salicifolius* chromosome 8 and on chromosome 8 itself, with an additional 12 found on a single X (Supplementary Table 4). Of these latter 12 genes, two (PRD1 and GYRA) have functions related to recombination, and another (ARF17) is involved in female germline specification. This suggests that some genes with functions related to sex chromosome evolution may have been recruited from this chromosome in particular species, but we cannot make strong claims about selection or functional enrichment from these data.

## Discussion

Chromosomal rearrangements are thought to play a pivotal role in evolution, promoting adaptation under a variety of circumstances. However, to date, phylogenetic approaches capable of explicitly testing these hypotheses in multispecies datasets have remained limited. Here, we have developed a new phylogenetic comparative approach to the analysis of multispecies synteny datasets, allowing for explicit hypothesis testing regarding the drivers of rearrangement. While our approach makes a number of assumptions, particularly regarding the statistical independence of adjacencies and presence/absence vs. count data, we believe the general framework presented here lays the groundwork for future advances, and our simulation approach to evaluating significance should enable more robust inferences about significant rate differences even if our assumptions are violated. Ultimately, we envision a statistical toolkit that can explicitly estimate rates of different kinds of chromosomal rearrangement and test for accelerated rates across species, phenotypes, and genomic regions, paving the way to a better understanding of the macroevolutionary drivers of genomic change.

We observed strikingly high rates of chromosomal rearrangement in *Rumex*, particularly in dioecious taxa. For sex chromosomes, this fluidity indicates extensive opportunities for gene movement towards and away from sex-linked regions. While past work in *R. hastatulus* revealed little evidence for evolutionary strata despite highly heterogeneous substitution rates between the X and Y chromosome (*21*), we can now understand that this reflects extremely high rates of rearrangement, leaving little spatial patterns of historical origins of sex-linked genes. The transition to dioecy from hermaphroditism, and accompanying evolution of genetic sex determination, could lead to elevated rates of syntenic evolution in at least two ways. First, sex chromosome inversions and X-autosome fusions can link sexually antagonistic variation, becoming favored to extend sex-linked regions (*3, 27*); evidence for accelerated intra-chromosomal rearrangements on the X in the XYY clade highlight this possible contribution. Second, the release of constraint on the expression of genes with sex-associated functions should lead to generally elevated rates of molecular evolution. Both factors may be at play in *Rumex*, though we cannot rule out the contributions of others factors associated with dioecy.

Mating system could influence rates of rearrangement independently of the effects of sexual antagonism. As obligate outcrossers, dioecious taxa should have lower linkage disequilibrium and therefore experience stronger selection for rearrangements as recombination modifiers; this could help explain why autosomal rearrangements also appear elevated among dioecious *Rumex* species, especially in the XY clade. Alternatively, underdominant rearrangements should spread faster in selfing populations (*28*); while the degree of selfing in our hermaphroditic species is unknown, our results appear inconsistent with this prediction. Outcrossers should also exhibit greater opportunities for meiotic drive and the activity of transposable elements than hermaphroditic taxa experiencing significant rates of selfing (29-30). Some of our dioecious taxa are also annual, with faster generation times potentially driving elevated rates of molecular evolution. However, broad increases in rates of molecular evolution cannot fully explain our results, as the degree of substitution rate variation we observe across coding sequences is much smaller than our estimated rate variation of syntenic changes (Supplementary Figure 16). In either case, these results highlight the possible importance of natural selection in driving accelerated rates of rearrangement in some taxa. In the future, it will be necessary to integrate our new methods and findings with population genomic approaches for inferring selection and demography to further understand what drives the remarkable rate of rearrangement we observe in this genus.

## Acknowledgements

We thank Thomas Gludovacz, Bill Cole, Olena Voznesenska, Alice DesRoches, Shawn Patille, and numerous work-study students for help with plant growth and maintenance. This work was supported by an NSERC Postdoctoral Fellowship awarded to M.S.H., startup funds provided to M.S.H. by the University of Rochester, NSERC Discovery Grants awarded to S.I.W and S.C.H.B., and an NSERC Canada Research Chair awarded to S.I.W. Source code and installation instructions for the *SynPhyMe* R package are available at https://github.com/mhibbins/SynPhyMe. Code and data related to manuscript analyses are available at https://github.com/mhibbins/rumex_synteny. Raw sequencing data and genome assemblies and annotations for *Rumex* will be uploaded and made available at GenBank.

