## Supplementary Materials and Methods for "Rapid evolution of synteny associated with multiple origins of dioecy and XY sex determination"

#### Assigning syntenic regions to phylogenetic characters

*SynPhyMe* takes two inputs: a csv file containing information on pairwise syntenic blocks phased to a reference genome and an ultrametric phylogenetic tree in newick format. The csv file must have a minimum of ten columns, where each row is an individual syntenic block shared by two species: genome1, genome2, chr1, chr2, startBp1, endBp1, startBp2, endBp2, refChr, and refGenome. Users may track custom genomic regions as defined in the reference genome using the useColors flag; in this case, the input file must include an additional "color" column, indicating the color assignment for each syntenic block. If useColors is false, each chromosome of the reference genome will be tracked as a region of interest. These regions of interest determine the resolution of syntenic regions and subsequent adjacencies to track. Each pairwise combination of regions (chromosomes or sub-chromosome genomic regions with color assignments) is a possible adjacency for which presence/absence can be mapped. This input file is formatted to match the "phasedBlks" output of the riparian plotting functions of GENESPACE (*15*), and these output files can be given directly to *SynPhyMe*.

Since syntenic blocks are estimated between pairs of genomes, we assign adjacencies to each species using a simple algorithm based on the comparison to their closest phylogenetic relative, with the goal of maximizing the resolution of synteny inference by minimizing evolutionary distance. Sister species are compared to each other; when the focal taxon is sister to an ingroup clade, the next earliest-diverging member of that clade is used for comparison; when this clade contains multiple taxa, one of them is sampled at random for comparison. Once a comparison is assigned, all syntenic blocks for the current species are sorted in order of physical position along each chromosome, and an adjacency is recorded any time the phased color or chromosome assignment shifts along a chromosome. Importantly, each adjacency is only recorded once as a presence/absence trait, so additional instances of the same adjacency are not counted.

These adjacencies are also unordered, such that the adjacencies *AB* and *BA* are treated as equivalent.

**Estimating evolutionary rates from adjacency data**

The Mk model in phylogenetics models the evolution of discrete characters using a matrix of transition probabilities between possible character states (*22-23*). In the simplest form of this model, a single rate parameter governs the rate of transition of 0 → 1 and 1 → 0 transitions for a binary character. This gives us a simple rate transition matrix, dubbed ***Q***:

$$\boldsymbol{Q} = \begin{matrix} -q & q \\ q & -q \end{matrix}$$

. Our goal is to estimate the value of $q$ over our presence/absence matrix of syntenic adjacencies. We do this using the pruning algorithm (*24*), which visits each node of the phylogeny in a postorder traversal, estimating ancestral state probabilities. For the Mk model, the transition probability matrix for a given branch is given by $P = e^{Qt}$, where $\boldsymbol{Q}$ is the transition rate matrix and $t$ is the length of the branch. Once character state probabilities are obtained at the root, a likelihood can be obtained by taking the product of these probabilities over some prior (in our case, a uniform prior).

Let $b = \{b_1, b_2, \dots\ b_n\}$ be the vector of color assignments or chromosomes phased to a reference genome in the input file. Then, let $A = \left\{\{b_i, b_j\} \mid 1 \ \leq i < j \ \leq n\right\}$ be the set of all unordered pairs (adjacencies) in ***b*** which are observed in at least one species. Our goal is to estimate a single value of $q$, describing overall rates of genomic rearrangement across all the elements of **A**. We do this by optimizing the following composite likelihood:

$$L = \sum_{A} \quad - log\ (L(A_i))$$

where $L(A_i)$ is the likelihood score obtained from the pruning algorithm for a single adjacency. Note that this metric is a pseudolikelihood, since the assumption of independence is violated when rearrangements affect multiple adjacencies. To avoid local optima, we propose ten initial guesses for the value of $q$, ranging from $10^{-5}$ to $10^{4}$. The value of $q$ is optimized on ***L*** for each initial guess using the Brent method with a lower bound of 0, using the *optim* function of the *stats* R package version 4.4.1. The maximum value of ***L*** across these ten optimizations is then chosen as the optimal value for *q*.

**Simulation study**

To assess the accuracy of our rate estimation approach, we conducted two simulation studies. In the first study, we randomly generated phylogenies of five taxa using the *rtree* function in the R package *ape* version 5.8 (*31*). For each of these phylogenies, we then simulated a matrix of 25 binary characters under the Mk model using the *simulate_mk_model* function in the R package *castor* version 1.8.2 (*32*). This process was repeated 300 times for each of 10 different simulated values of $q$, ranging from 0.01 to 0.1, and the composite likelihood across the 25 binary characters was optimized to estimate the value of $q$ for each replicate.

For the second study, we implemented a new forward-time phylogenetic simulator of chromosomal rearrangements in our R package *SynPhyMe*. Users specify the phylogeny, the initial karyotype at the root, and the chromosomal mutation rate. The initial karyotype is a dataframe with three columns: chromosome, color, and position, indicating the initial chromosome configuration and the positions and color IDs of syntenic blocks to track through the simulation. Each branch of the phylogeny is visited from the root to the tips in a preorder traversal. On each branch, a number of chromosomal mutations is drawn from the Poisson distribution with $\lambda = \mu t$, where $\mu$ is the user-specified chromosomal mutation rate per unit time on the phylogeny, and $t$ is the length of the current branch. If at least one mutation is drawn, a rearrangement type is randomly chosen from a list including chromosomal fusions and fissions, inversions, and reciprocal and nonreciprocal translocations. Users may also specify their own rearrangement list containing a subset of these types. When a mutation occurs on an ancestral node, the rearrangement is inherited by all descendants of that node.

When implementing rearrangements, we assume holocentric chromosomes, such that rearrangement breakpoints can occur anywhere along a chromosome. For fusions, we randomly sample two chromosomes and join them at the ends. For fissions, we randomly select a chromosome and then randomly choose a breakpoint position on that chromosome to split. For inversions, we randomly choose a chromosome, randomly choose two breakpoint positions on that chromosome, and reverse the positional order of syntenic color assignments within the bounds of those breakpoints. For translocations, two chromosomes are randomly chosen; for reciprocal translocations, a breakpoint is chosen at random on each chromosome, and the ends of the chromosomes from the breakpoint onward are swapped. For nonreciprocal translocations, a breakpoint is chosen on one chromosome, and the end of that chromosome from the breakpoint onward is moved to the end of the other chromosome.

The output of the simulation is the same three-column dataframe containing the updated chromosomes and syntenic regions for each species. These can be converted into adjacencies mapped on the specified phylogeny by *SynPhyMe* for use in downstream analyses. For our study, we simulated an initial karyotype of 10 chromosomes, each broken into three syntenic regions, and randomly simulated five-taxon trees using *rtree*. We conducted 300 replicate karyotype simulations for each of ten values of the chromosomal mutation rate, ranging from 0.1 to 1. Each simulated karyotype was mapped to the simulated phylogeny as an adjacency matrix, and we used our composite likelihood approach to estimate $q$ values.

**Testing for shifts in the rate of rearrangement**

To allow for hypothesis testing relating to rates of rearrangement, we implemented a simple two-rate version of the Mk model. For this analysis, users specify a list of focal input taxa that are a subset of those in the phylogeny. For these species, an additional scaling parameter called $r$ is introduced, which serves as a rate multiplier for the transition rate matrix, $\boldsymbol{Q}$. When the pruning algorithm visits a branch on the phylogeny that is ancestral to a monophyletic clade of focal taxa, or a leaf corresponding to a focal taxon, the following matrix is used to calculate transition probabilities:

$$Qr = \ [-qr\, qr\, qr\ \ -qr\,]$$

For other branches, the standard transition rate matrix $\boldsymbol{Q}$ is used (*33-34*). The one-rate model is a special case of the two-rate model where $r = 1$, so these models are nested. To avoid local optima, we optimize over a 10x10 grid of initial $q$ values ranging from $10^{-5}$ to $10^{4}$ and $r$ values ranging from 0.1 to 10. Optimization is done with the *optim* function in the R package *stats*, using the L-BFGS-B method with a lower bound of 0.

Our composite likelihood approach violates the assumptions of standard methods for assessing the fit of nested models, like AIC and BIC. Therefore, we implemented a simulation-based approach to evaluate the significance of the two-rate model. After $q$ is estimated on the data for the one-rate model, we simulate a large number of replicate binary character datasets under the one-rate model using the empirical phylogeny, length of the empirical adjacency matrix, and the estimated value of $q$. For each of these simulated datasets, we estimate $q$ under the one-rate model, as well as $q$ and $r$ under the two-rate model. If $L_1$ and $L_2$ are the optimal likelihood scores for the one-rate and two-rate models, respectively, we calculate the likelihood-ratio test score as

$$LRT = 2[abs|-\, log\; log\; (L_1)\; | - abs| - log\; (L_2)|]$$

Applied to each replicate simulation, we generate a null distribution of LRT scores. The LRT score obtained from the empirical one and two-rate models is assigned rank $i$ against this distribution, where $i = n + 1$ indicates the empirical LRT is larger than all values in the distribution of null LRTs. Finally, the p-value for the two-rate model is calculated as

$$p = 1 - \left(\frac{i-1}{n}\right)$$

where $n$ is the number of replicates in the simulated distribution.

#### *Rumex* genome assembly

For the genome assemblies, individuals were grown in the glasshouse from multiple collection trips. In 2017, *Rumex paucifoliu*s was collected from Teton County Wyoming. *Rumex scutatus* and *R. sagittatus* were collected from Tenerife, Canary Islands and Tsitsikamma National Park, South Africa, respectively. We also used seed from the Kew Millenium Seed Bank from Greece for the genome assembly of *R. acetosa*, seed collected from the Botanical Garden in Tel Aviv, Israel for *R. rothschildianus*, and seed from the China Southwest Genobank for *R. acetosella*. All seeds were germinated on moist filter paper in a petri dish at 4 degrees Celsius for 20 days then at 22 degrees Celsius 16-hour day-length for 7 days. The seeds were then transplanted to 0.5L pots containing a mix of 1:1 sunshine mix and profile. Glasshouse temperatures were between 20 and 24 degrees Celsius with a 16-hour daylength and watered as needed. Plants were fertilized once per week beginning two weeks after planting with a 20-20-20 fertilizer. Plants were transferred to bigger pots as they became pot-bound.

We used our recently assembled genomes of *R. salicifolius*, the XYY maternal cytotype of *R. hastatulus* (*21*) and the XY maternal cytotype of *R. hastatulus* (*26*). We also used a recent genome assembly for *R. bucephalophorus* (*18*). For the remaining species we generated long-read de novo assemblies using HiFi PAC Bio sequencing and Dovetail OmniC sequencing. All sequencing was completed at Dovetail Genomics except for *R. acetosa*, which was sequenced at Hudson Alpha. For all remaining species assemblies, we used hifiasm v0.19.6 (*35*) to assemble the genomes using default parameters into phased haplotypes for both dioecious and hermaphroditic species. Paired-end reads were then mapped to the resulting haplotype assemblies using bwa v0.7.17 (*36*) and filtered following the Arima mapping pipeline with a MapQ > 12 (https://github.com/ArimaGenomics/mapping_pipeline). The resulting bam files had duplicates removed using Picard v2.7.1 (https://broadinstitute.github.io/picard/). We then scaffolded both haplotypes using YaHS v1.2a.2 using default parameters (*37*). Using plots of synteny between both haplotypes of dioecious species as well as contact maps for all assemblies, we visually inspected each chromosome for miss-assemblies or potentially heterozygous copies (see below for details).

For the hermaphroditic species, both *R. sagittatus* and *R. scutatus*, we used the first haplotype for all analyses, as both haplotypes for each species were complementary. We compared both haplotype of each species using dot plots generated by genespace analyses (see below for details). For *R. acetosella*, we similarly used one of the two haplotypes for the remaining analyses. Although *R. acetosella* is a dioecious species with an XY sex chromosome system, we could not determine the location of the sex chromosomes; therefore, remaining analyses will only use the 9 largest scaffolds in the assembly. For the remaining dioecious species we used a combination of sequencing data, HiC contact maps, and syntenic analyses to detect sex-linked regions of the genome and potential miss-assemblies.

For the XYY species, *R. rothschildianus* and *R. acetosa*, we detected sex chromosomes using genomic coverage between sexes. We used whole genome sequencing data from one male and one female individual for both assemblies. For *R. rothschildianus* these data were generated by Genome Quebec while the *R. acetosa* data were accessed via NCBI's SRA database (Bioproject accession PRJNA914205). These data were whole genome sequencing data from individuals from Greece. Using these data, we mapped them to each haplotype for each assembly using bwa v0.7.17 (*36*). Resulting bam files were sorted by genomic coordinate, filtered by a mapping quality of at least 30, and duplicates marked using Picard v2.7.1 (https://broadinstitute.github.io/picard/). Coverage was quantified and compared using tinycov v0.4.0 (https://github.com/cmdoret/tinycov). Scaffolds that had similar coverage between sexes are most likely autosomes, while scaffolds that have higher coverage in females are most likely X-linked. Lastly, scaffolds that have higher coverage in males are most likely Y-linked. We then generated contact maps using the Dovetail Linked-Read Analysis pipeline (https://dovetail-analysis.readthedocs.io/en/latest/data_processing/contact_map.html). First, PacBio reads were aligned to the assemblies using bwa v0.7.17. Pairtools v1.1.2 (*38*) was then used to create a pairsam file containing potential pairs using a minimum threshold of 40 for defining multimapping alignments and maximum gap of 30 between alignments. We then used pairtools sorted, dedup, and split to sort the resulting pairsam, mark duplicate pairs, and split into two files. Lastly, the pairs file was then used to generate the final HiC files using juicer v1.22.01 (*39*). The contact map for the phased maternal (X-bearing) haplotype was visually inspected in Juicebox to identify false joins and other assembly errors (*40*). In particular for *R. rothschildianus*, breaks were inserted into the original scaffold 1 to create chromosomes 1, 3, 4, 5 and X. Chromosomes 3 and 5 were then reverse complemented to resolve mis-assembly errors. Chromosome 2 was unchanged from scaffold 2 and chromosome 6 was renamed from scaffold 4. The resulting assembly is the X bearing haplotype which was used for all future analyses. The Y bearing haplotype contains all autosomes and Y1 and Y2 which were assembled separately and required no adjustments. For *R. acetosa*, we resolved a false join in scaffold 3 by breaking the scaffold into two pieces. Both X-bearing assemblies had 7 scaffolds (6 autosomes and the X), corresponding to the known chromosome number for *R. rothschildianus* and *R. acetosa* (*41-42*). The Y bearing haplotype for *R. acetosa* contained a large Y scaffold with smaller Y-linked scaffolds rather than separately assembled Y1 and Y2 chromosomes.

For the XY species, *R. paucifolius,* we used a genomic coverage analysis to detect sex-linked regions. We used PacBio sequencing data of 2 males and 2 females to detect sex-linked regions. These data were generated by Sick Kids. We mapped these samples to each of the scaffolded haplotypes using pbmm2, a C++ wrapper for minimap2. The resulting bam files were then sorted by genomic coordinate and filtered by a quality score of 30 using samtools (*43*) and duplicates marked with Picard v2.7.1 (https://broadinstitute.github.io/picard/). We then quantified coverage across the genome for the males and females using tinycov v0.4.0. Similar to analyses for the XYY species, scaffolds with variation in coverage between sexes are most likely sex-linked. For *R. paucifolius*, we also used male to female coverage ratios to detect sex-linked regions. After identifying the sex chromosomes, we used our generated contact maps to resolve mis-assemblies. For the maternal haplotype had a false join in scaffold 2. Therefore, using syntenic hits between both haplotypes and the agp file from YAHS, we split scaffold 2 into two scaffolds. For the paternal haplotype, there was a false join on scaffold 1 that joined autosomal scaffold 1 and the Y chromosome.

#### ***Rumex* genome annotation**

The *Rumex* genome annotations were created using a combination of MAKER-3.01.03 (*44*) and liftoff (*45*). The Maker annotation of *R. hastatulus* (TX haplotype), *R. acetosa*, *R. rothschildianus*, *R. sagittatus*, *R. paucifolius*, and *R. scutatus* genomes was performed as described in (*21*), except that, de novo repeats were annotated using RepeatModeler (*46*) for *R. hastatulus*, *R. acetosa* and *R. rothschildianus*.

We used previously published genome annotations for the XYY cytotype of *Rumex hastatulus* and *R. salicifolius* (*21*). This *R. hastatulus* annotation and corresponding genome assembly were used to create a genome annotation for *R. acetosella* using liftoff v1.6.3. This annotation was polished using the finalized version of one of the haplotype genome assemblies using the following alignment parameters: -a of 0.8, -s of 0.8, and -flank of 1. For the *R. bucephalophoru*s annotation we used the 8 largest scaffolds of the published genome assembly and its corresponding annotation to lift off an annotation with just 8 scaffolds using default parameters. For *R. paucifoliu*s, we used liftoff to create an annotation without Y linked scaffolds.

**Inference of *Rumex* syntenic regions and phylogenetic relationships**

To characterize patterns of synteny between protein-coding genes in the *Rumex* hermaphroditic and dioecious species we used the R package GENESPACE v1.3.1 with default parameters (*15*). For these and later analyses, we chose *Rumex salicifolius* as the outgroup species, as it belongs to a clade which is likely ancestrally hermaphroditic and has a more stable and likely ancestral karyotype (*16-17*). For each species, we used the corresponding annotation as described above and a peptide file generated using gffread v0.12.3 (*47*). GENESPACE uses Orthofinder v2.5.4 (*48*) and MCScanX (*49*) to scan synthetic blocks for orthologs and paralogs.

To construct an updated phylogeny of *Rumex* using our new genome assemblies, we queried the pangene output of our GENESPACE run to generate a pangenome annotation file. From this file, we extracted a list of 2732 single-copy syntenic orthologs with copies present in all 10 genomes for downstream phylogenetic inference. Using the annotation nucleotide and peptide sequences for these orthologs, we inferred peptide alignments using MUSCLE v3.8.31 (*50*) and codon alignments using RevTrans (*51*). Codon alignments were concatenated to create a single alignment of 5.5 Mb, which was then filtered using Gblocks (*52*) for codon sequences with the -b5 setting allowing for gap positions, resulting in a filtered alignment of 3.9 Mb. This alignment was given to IQ-TREE v.2.1.4 (*53*) for phylogeny estimation using ModelFinder, SH-aLRT, and ultrafast bootstrap with 1000 replicates.

To time-calibrate our maximum-likelihood phylogeny for downstream analyses of synteny evolution, we used a penalized likelihood approach implemented in the *chronos* function of the R package *ape* (*54*). We constrained the root of *Rumex* to a maximum age of 23 MYA based on fossil evidence (*55, 56*), the split of the two cytotypes of *R. hastatulus* to 0.8 MYA based on demographic modelling (*20*), and the split of *R. bucephalophorus* from the XY clade to 10 MYA based on a previous time calibration (*18*). We evaluated correlated, discrete, and relaxed clock models, choosing the model with the lowest PHIIC score.

**Estimation of rates of genomic rearrangement in *Rumex***

We used our two-rate approach implemented in *SynPhyMe* to test if the dioecious *Rumex* species in our dataset (*R. acetosa*, *R. rothschildianus*, *R. acetosella*, *R. paucifolius*, and the two cytotypes of *R. hastatulus*) exhibit faster rates of genomic rearrangement than their hermaphroditic relatives. For the syntenic blocks dataset, we used the phased blocks output of our GENESPACE analysis with *R. salicifolius* as the reference genome, and each of the ten chromosomes of *R. salicifolius* as tracked syntenic regions, resulting in 45 possible adjacencies. To avoid over-counting syntenic adjacencies on the homologous regions of X and Y chromosomes, we excluded Y chromosomes from this analysis. We used our time-calibrated phylogeny estimated from syntenic orthologs as the input phylogeny.

After converting syntenic regions to adjacency characters, we obtained a binary presence-absence matrix of 28 adjacencies observed in at least one of ten sampled species. Applying our composite likelihood approach to these data, we estimated a Q value of 0.022 under the one-rate model. To generate a null distribution of likelihood-ratio test scores for evaluating the significance of the two-rate model, we then simulated 100 datasets of 28 binary characters each under our estimated Q value using the *Rumex* phylogeny. The LRT value obtained from our two-rate analysis was ranked against this simulated distribution to evaluate significance of the model fit.

**Syntenic regions associated with sex chromosome evolution**

To test if particular syntenic adjacencies are correlated with the evolution of dioecy across our phylogeny, we fit phylogenetic generalized linear mixed models (PGLMMs) using the R package *phylolm* version 2.6.5 (*57*). For each adjacency independently, we fit a logistic model with sexual system ("hermaphroditic" or "dioecious") as a fixed effect and our estimated phylogeny as a random effect. We used flat priors for the fixed effects, and a fixed residual variance to model categorical data. We used the "logistic_MPLE" method to fit each model, which maximises the penalized likelihood of the regression. To evaluate the statistical power of this analysis, we also fit a model for a hypothetical "perfect" binary trait which is present in all of the dioecious species and absent in all hermaphroditic species. We found a weakly significant ($p = 0.0385$) relation between sexual system and this hypothetical trait, indicating limited power of this analysis to detect significant associations. For this reason, we interpret the results of this analysis with some caution.

We found a marginally significant ($p < 0.1$) association of three syntenic adjacencies with sexual system: *Scaffold_5Scaffold_8*, *Scaffold8_Scaffold7*, and *Scaffold_3Scaffold_8*. All three of these adjacencies involved chromosome 8 of *R. salicifolius*, so we flagged this chromosome for follow-up analyses to identify potential genes or regions with sex-associated functions. Parsing the phased blocks file from our GENESPACE run, we found that chromosome 8 was the only *R. salicifolius* chromosome with syntenic regions present on all X chromosomes in our dataset (Supplementary Figure 1). To test if genes from this chromosome are significantly enriched on the X chromosomes, we conducted a permutation analysis. We randomly sampled sets of 2769 genes (the number of genes found on chromosome 8) from the genome annotation of *R. salicifolius*, and calculated proportion overlap scores of these gene sets against the set of genes found on any X chromosome in our dataset. We then ranked the empirical overlap of chromosome 8 genes with X genes against this distribution, and evaluated significance using the formula described earlier under *Testing for shifts in the rate of rearrangement*. We found no significant enrichment of chromosome 8 genes on the sex chromosomes ($p = 0.1732$).

Finally, we used the pangene files and phased block coordinates provided by GENESPACE to identify genes present in regions of the X chromosomes that are syntenic to *R. salicifolius* chromosome 8, and where copies are still present in both species. This returned a list of 14 genes, which is too small a set to have sufficient statistical power for GO enrichment analyses. We instead manually curated functional annotation information for each of these genes (Supplementary Table 4).

### Results

**Genome sequencing and assembly**

We assembled phased genome assemblies for seven species including both dioecious and hermaphroditic species. The maternal haplotype of the XY cytotype of *R. hastatulus* had a final assembly size of 1.497 Gb while the paternal haplotype was 1.863 Gb. 96% and 91% of reads assembled in five main scaffolds, respectively. These five scaffolds (4 autosomes, 1 sex chromosome) correspond to the expected chromosome number in the XY cytotype of *R. hastatulus*. The maternal haplotype had a BUSCO completeness score of 97.7% (Eukaryota database) and 94.3% (Embryophyta). The paternal haplotype had a BUSCO completeness score of 99.2% (Eukaryota database) and 95.1% (Embryophyta database).

Closely related species, *R. paucifolius*, had a genome assembly size of 908.7 Mb for the maternal haplotype and 932.9 Mb for the paternal haplotype. After breaking the false joins on each of the haplotypes, 94% and 97% of the maternal and paternal haplotypes respectively, assembled into the main 7 scaffolds, which corresponds to the expected chromosome number of *R. paucifolius.* The maternal haplotype had a BUSCO score of 100% (Eukaryota database) and 96.3% (Embryophyta databases). The paternal haplotype had a BUSCO completeness score of 100% (Eukaryota database) and 96.2% (Embryophyta databases).

Haplotype 1 of the first haplotype of *R. acetosella* was 2.489 Gb while the second was 1.418 Gb. Although the individual used for our reference genome is tetraploid, we assembled into two haplotypes for ease of analysis. The difference in assembly size is most likely due to repeat elements in the first haplotype due to its polyploid nature. 40% and 62% of reads, respectively, assembled into 7 scaffolds, which corresponds to the expected chromosome number. Haplotype 1 had a BUSCO completeness score of 100% (Eukaryota) and 97.8% (Embryophyta). The second haplotype had a BUSCO completeness score of 97.7% (Eukaryota) and 94.8% (Embryophyta).

The XYY species in our dataset, *R. acetosa* and *R. rothschildianus,* assembled well with the phased haplotypes representing the maternal and paternal haplotypes. The maternal haplotype of *R. acetosa* had an assembly size of 2.874 Gb while the second was 2.990 Gb. After breaking a false join, 73% of reads in the maternal haplotype assembled into 7 scaffolds (6 autosomes, 1 X chromosome). The maternal haplotype had a BUSCO completeness score of 97.7% (Eukaryota) and 96.6% (Embryophyta). 75% of reads in the paternal haplotype assembled into 6 autosomal scaffolds, with most of the remaining reads assembling into many small Y-linked scaffolds. The paternal haplotype had a BUSCO completeness score of 97.7% (Eukaryota) and 87.1% (Embryophyta).

The final assembly size of haplotype 1 of *R. rothschildianus* is 3.081 Gb while haplotype 2 is 2.976 Gb. Before the rearrangement of sex chromosomes, 69% of reads in haplotype 1 assembled into 7 chromosomes (6 autosomes, 1 sex chromosome) and 92% of reads of haplotype 2 assembled into 6 autosomes. Y1 and Y2 were originally assembled in haplotype 1 and were moved into haplotype 2, creating the maternal and paternal haplotypes. The maternal haplotype had a BUSCO completeness score of 98.4% (Eukaryota) and 96.4% (Embryophyta). The paternal haplotype had a BUSCO completeness score of 98.4% (Eukaryota) and 85.5% (Embryophyta).

For the hermaphroditic species, the first haplotype of *R. sagittatus* was 1.330 Gb and 2.560 Gb for *R. scutatus*. Only the first haplotypes were used in the following analyses for the hermaphroditic species. 96% of reads assembled into 9 scaffolds in *R. sagittatus*, corresponding to the expected chromosome number. *R. sagittatus* had a BUSCO completeness score of 100% (Eukaryota) and 96.2% (Embryophyta). Although the expected chromosome number in *R. scutatus* is 10, 96% of reads assembled into 9 scaffolds with the remaining being very small scaffolds. *R. scutatus* had a BUSCO completeness score of 99.2% (Eukaryota) and 96.7% (Embryophyta).

**Detection of sex-linked regions**

Using a genomic coverage analysis, we found regions of the genome in *R. acetosa* with differential coverage in males and females. This variation in coverage corresponds to expected coverage differences in sex chromosomes. In the maternal haplotype, coverage on the chromosomes in males was approximately double than in females, with the exception of one chromosome which had the same coverage in males and females (Supplemental Figure 4). Since females have two X chromosomes to the one in males, we expect coverage in females to be double on the X chromosomes. Considering all chromosomes had half coverage in females, except for the one that had equal coverage in both sexes, we deduced that this chromosome was the X. In the paternal haplotype, we saw a similar pattern, where most chromosomes had half coverage in females (Supplemental Figures 5). Additionally, many small scaffolds had coverage in males with little to no coverage in females, which is what we would expect for Y-linked regions. Therefore, the paternal haplotype has many small pieces of the Y1 and Y2 chromosomes. We see very similar patterns in *R. rothschildianus*, where coverage on the X in the maternal haplotype is approximately double in females (Supplemental Figure 2). In the paternal haplotype, we see little to no coverage on Y1 and Y2 in the female, with higher coverage in the male (Supplemental Figure 3). With the final *R. rothschildianus* assemblies, we detected 5,564 sex-linked SNPs in the maternal haplotype with 99% of the sex-linked SNPs on the X chromosome. We also detected 4,286 sex-linked SNPs in the paternal haplotype with 92% on Y1 and Y2. Using the *R. paucifolius* genome assemblies and PacBio sequences for males and females, we detected sex-linked regions associated with the X and Y chromosomes. We found a region with reduced male to female coverage ratio on the maternal haplotype which we termed the X chromosome (Supplemental Figure 6) and very high male to female coverage ratio on the paternal haplotype which we termed the Y (Supplemental Figure 7).

### Supplemental Figures and Tables

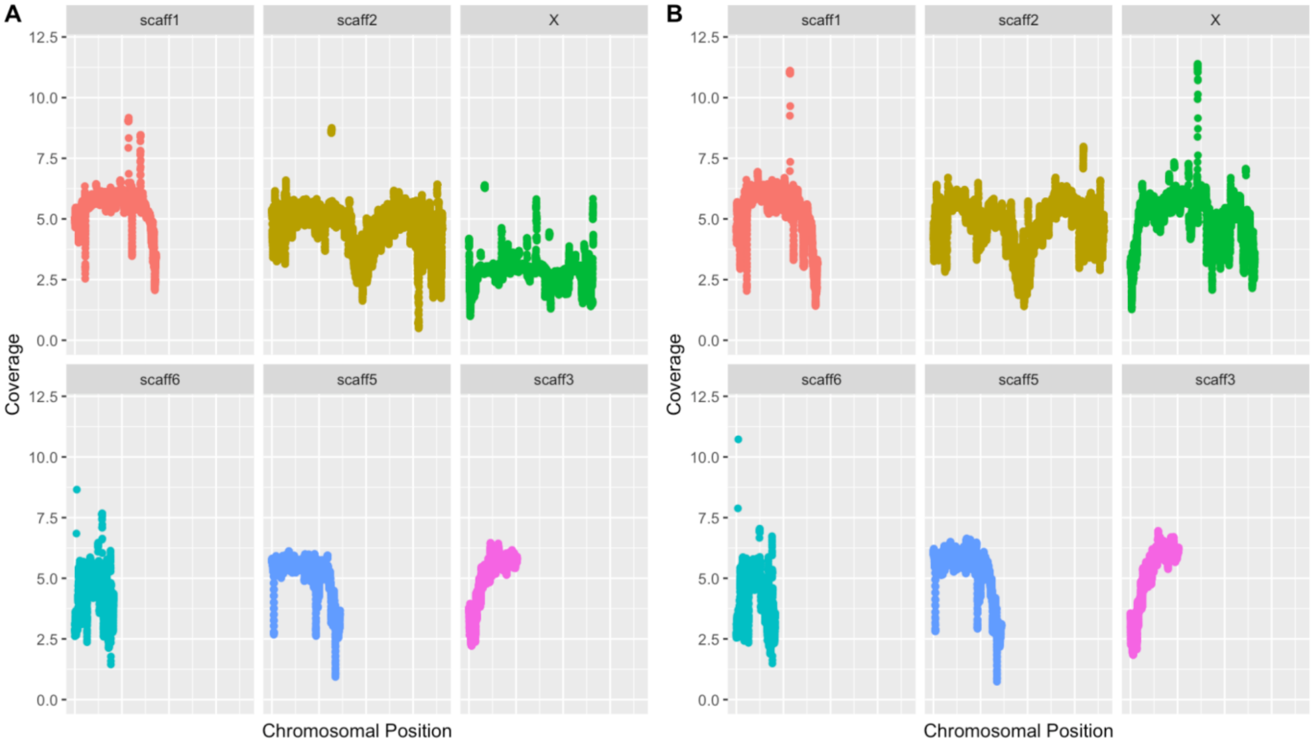


***Supplementary Figure 1.*** Genomic coverage across A. male genome and B. female genome using the maternal haplotype of *Rumex rothschildianus.* 5 autosomal scaffolds and the X chromosome are plotted, showing higher coverage on the X chromosome in females compared to males.

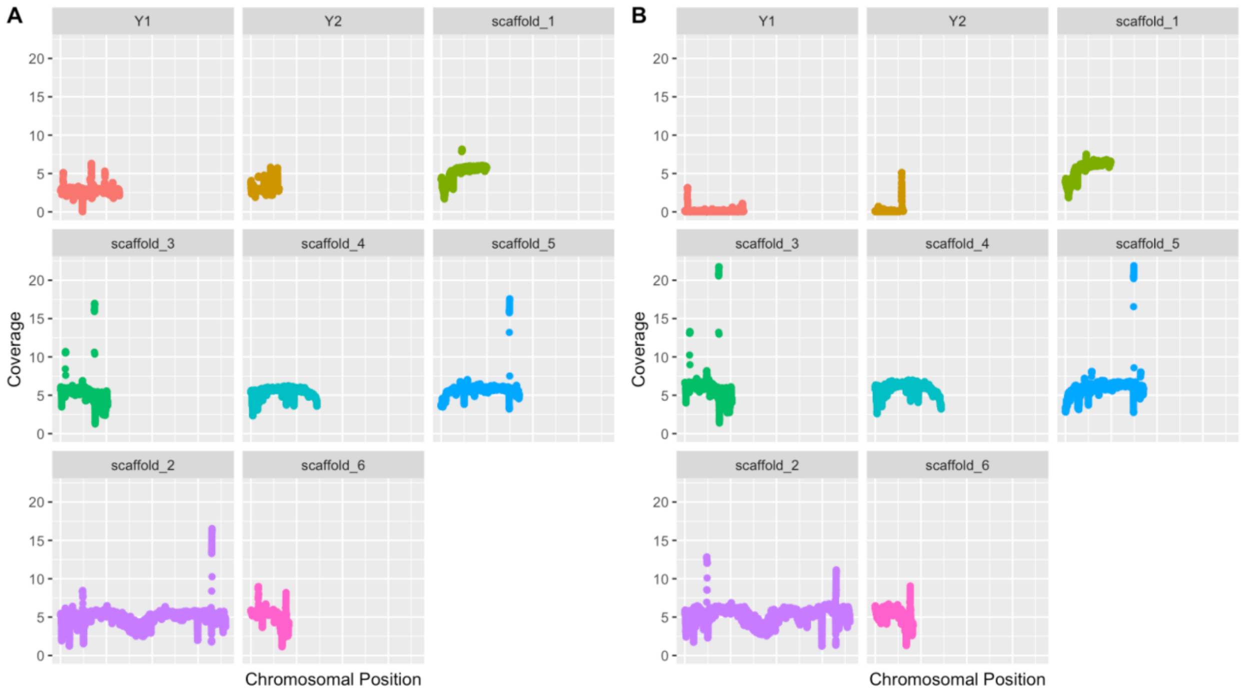


***Supplementary Figure 2.*** Genomic coverage across A. male genome and B. female genome using the paternal haplotype of *Rumex rothschildianus.* 6 autosomal scaffolds and both Y chromosomes are plotted.

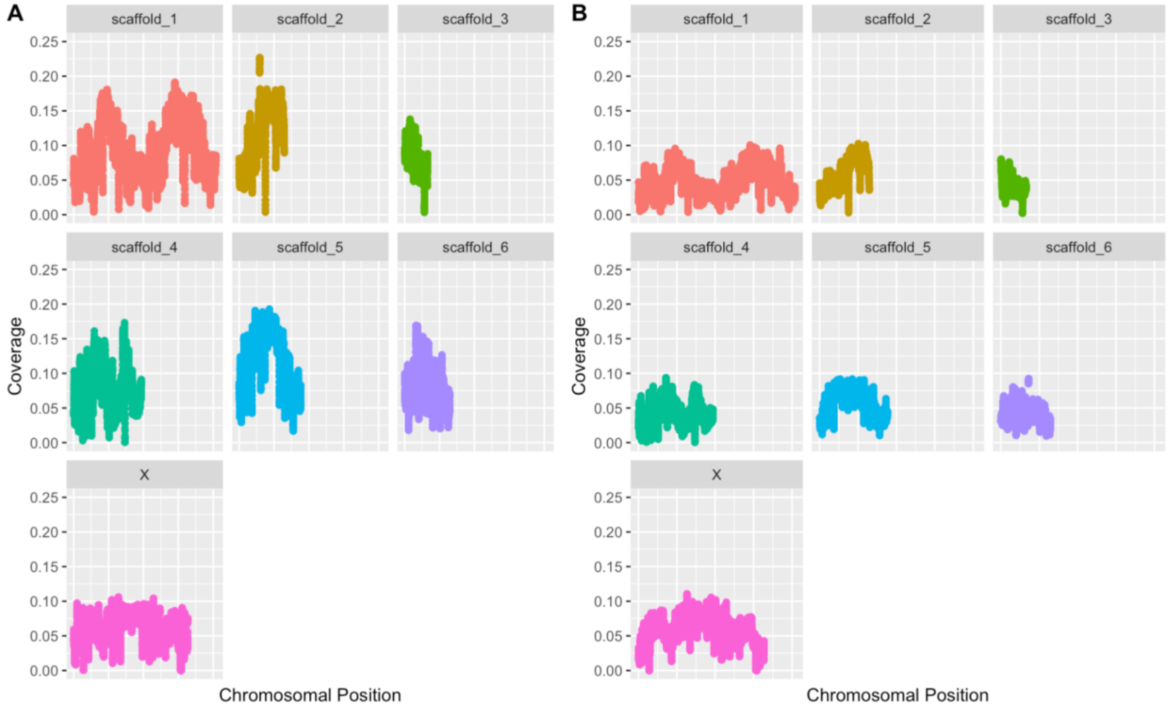

***Supplementary Figure 3.*** Genomic coverage across A. male genome and B. female genome using the maternal haplotype of *Rumex acetosa.* 6 autosomal scaffolds and the X chromosome are plotted.

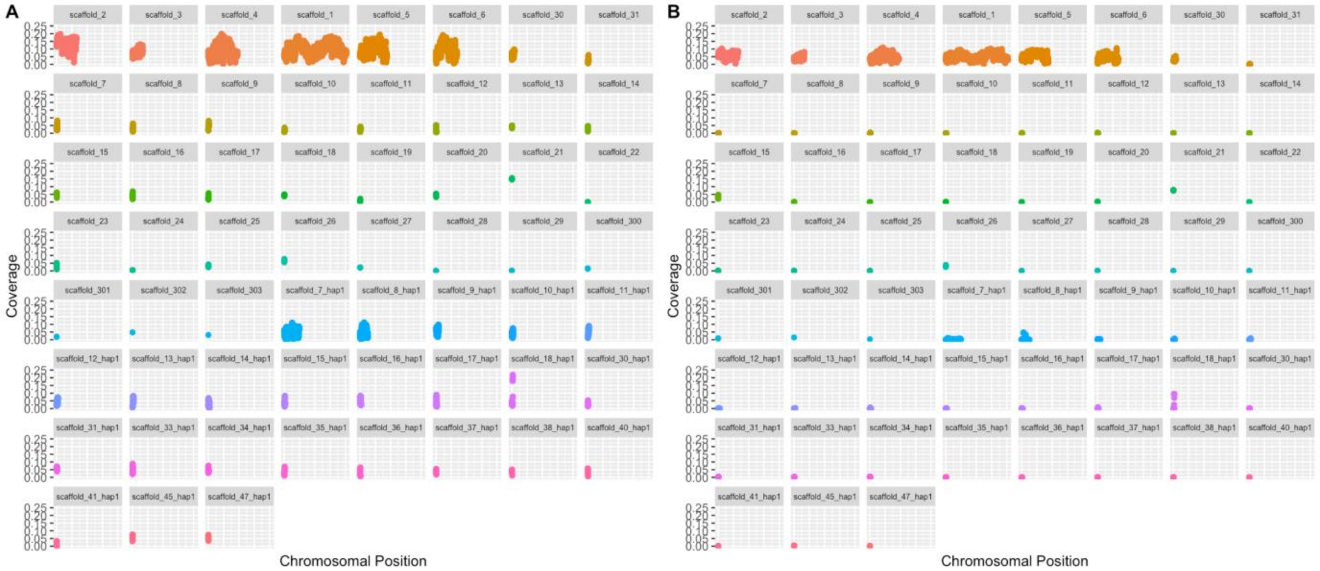


***Supplementary Figure 4.*** Genomic coverage across A. male genome and B. female genome using the paternal haplotype of *Rumex acetosa.* Scaffolds 1-6 are the autosomal scaffolds, while the remaining scaffolds are all small pieces of the sex-linked region or pseudo-autosomal region of the Y chromosomes.

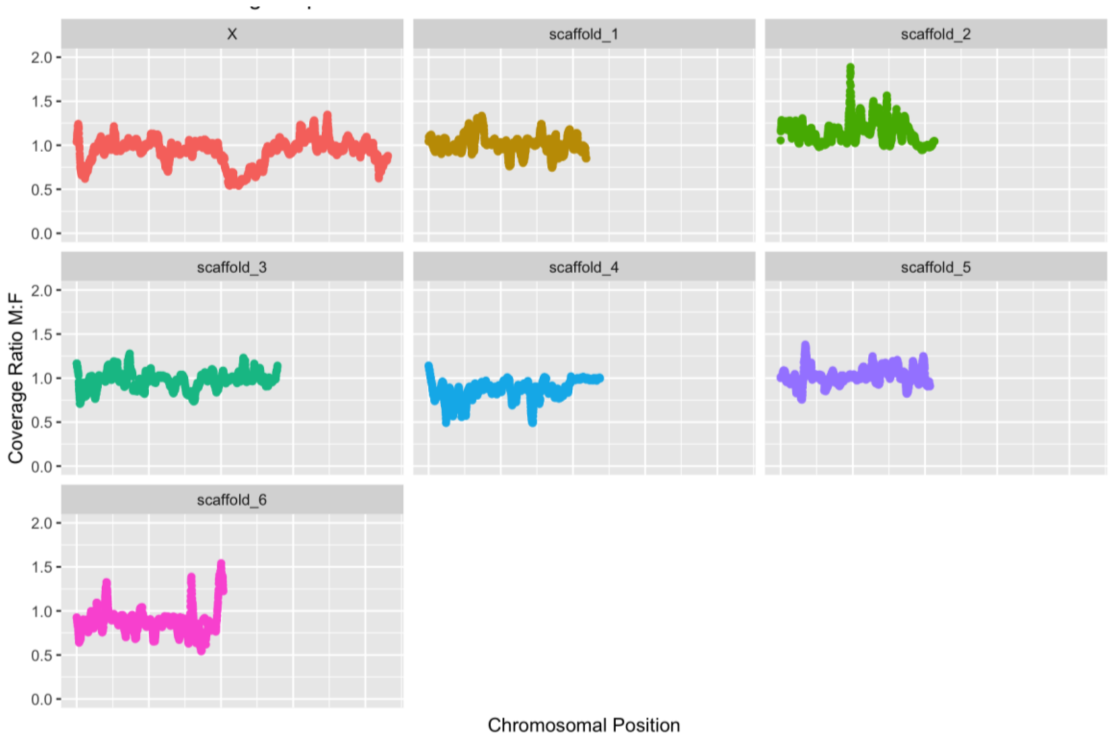

***Supplementary Figure 5:*** Male to female genomic coverage ratio across maternal haplotype of *Rumex paucifolius.* 6 autosomal scaffolds and the X chromosome are plotted, showing the dip in M:F coverage ratio on the X chromosome.

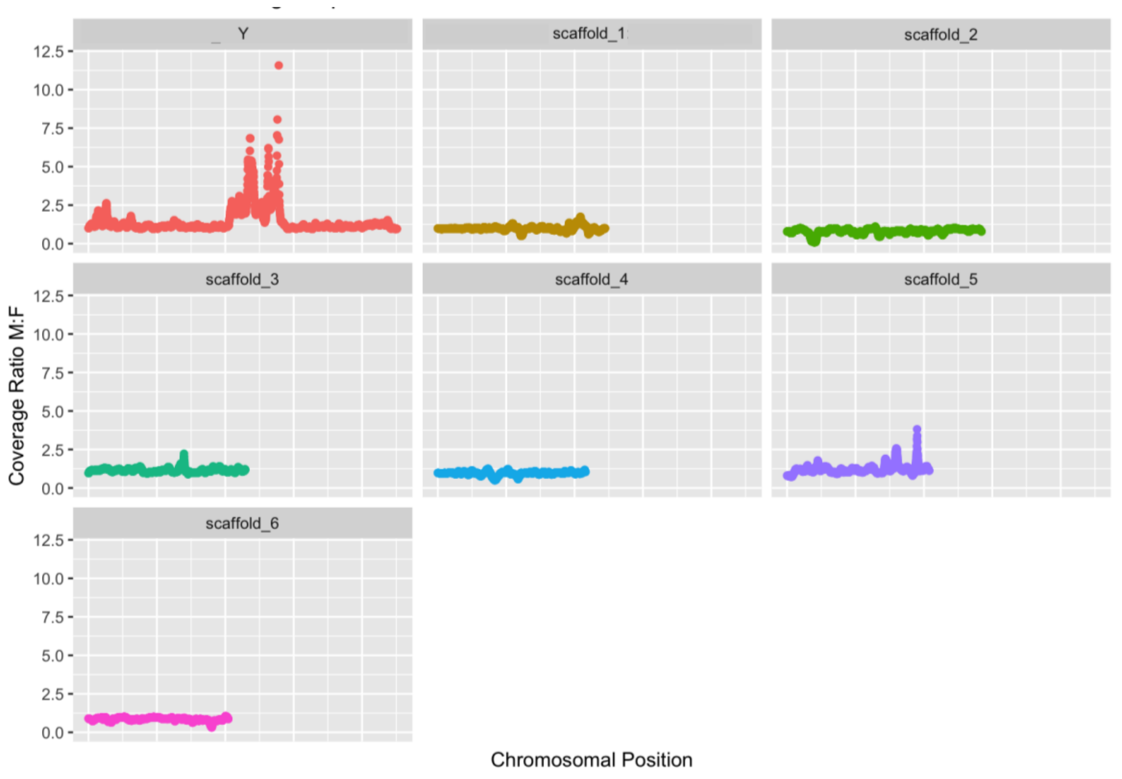


***Supplementary Figure 6.*** Male to female genomic coverage ratio across paternal haplotype of *Rumex paucifolius.* 6 autosomal scaffolds and the Y chromosome are plotted, showing a large increase in M:F coverage ratio on the Y chromosome.

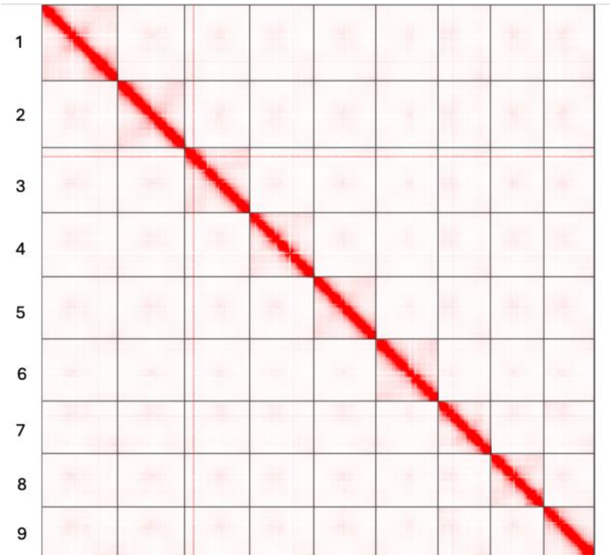


***Supplementary Figure 7.*** Omni-C contact map for *R. scutatus* assembly.

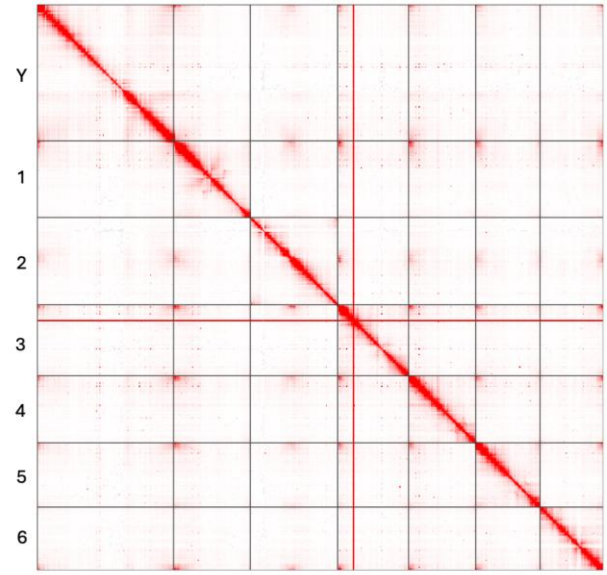

***Supplementary Figure 8.*** Omni-C contact map for haplotype 2 of *R. paucifolius* assembly.

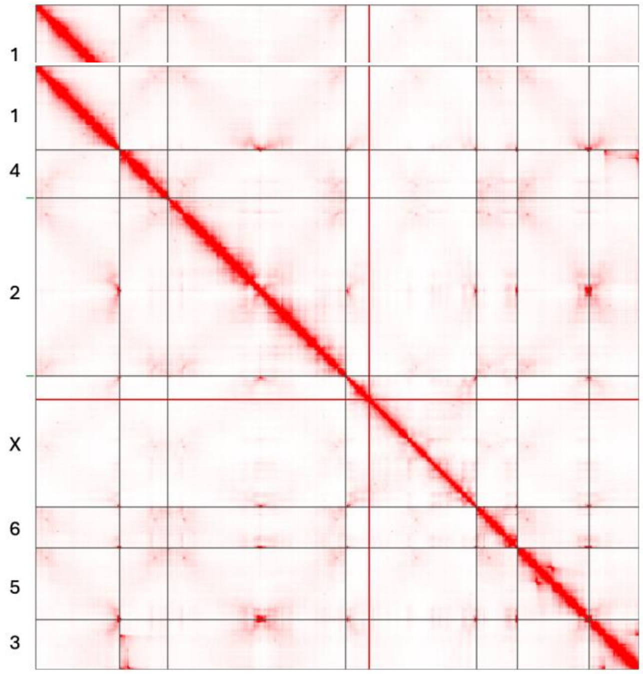


***Supplementary Figure 9.*** Omni-C contact map for haplotype 1 of *R. rothschildianus* assembly.

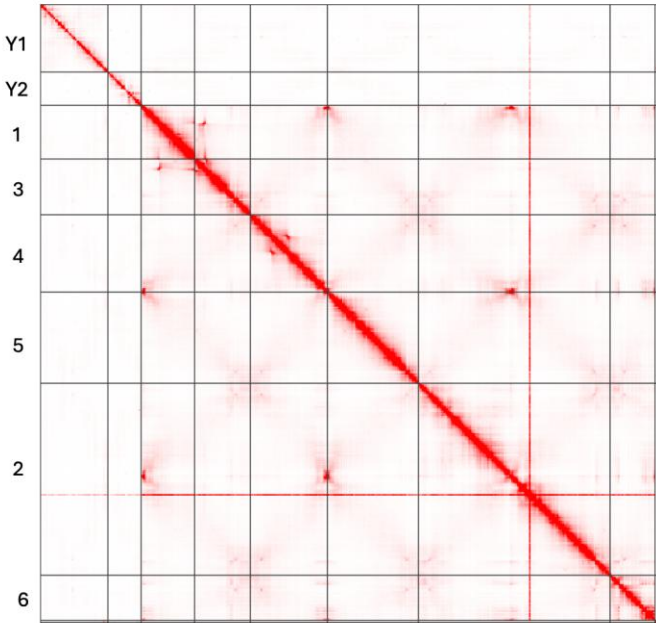


***Supplementary Figure 10.*** Omni-C contact map for haplotype 2 of *R. rothschildianus* assembly.

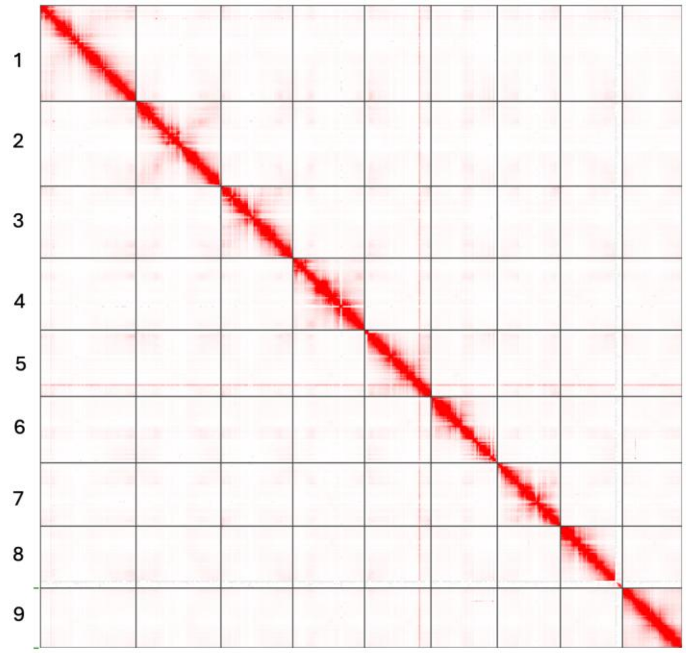


***Supplementary Figure 11.*** Omni-C contact map for *R. sagittatus* assembly.

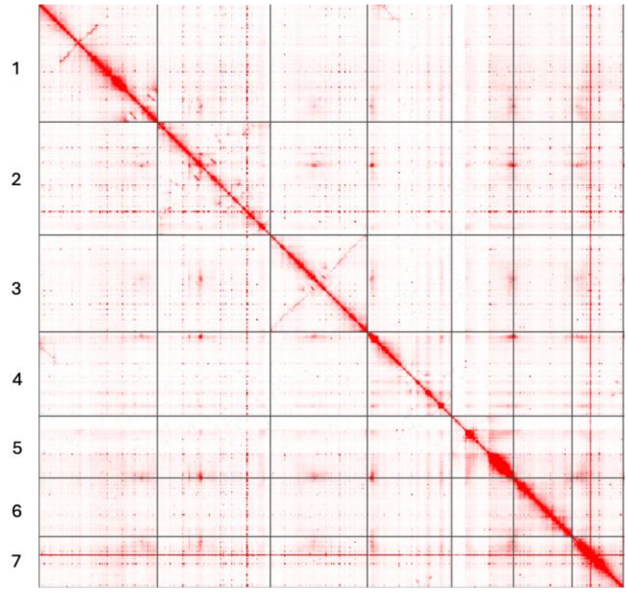


***Supplementary Figure 12.*** Omni-C contact map for *R. acetosella* assembly.

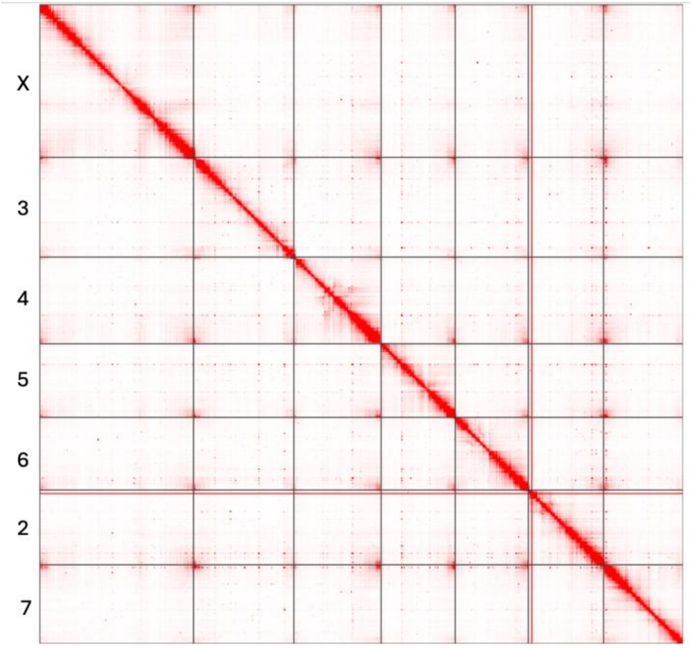


***Supplementary Figure 13.*** Omni-C contact map for haplotype 1 of *R. paucifolius* assembly.

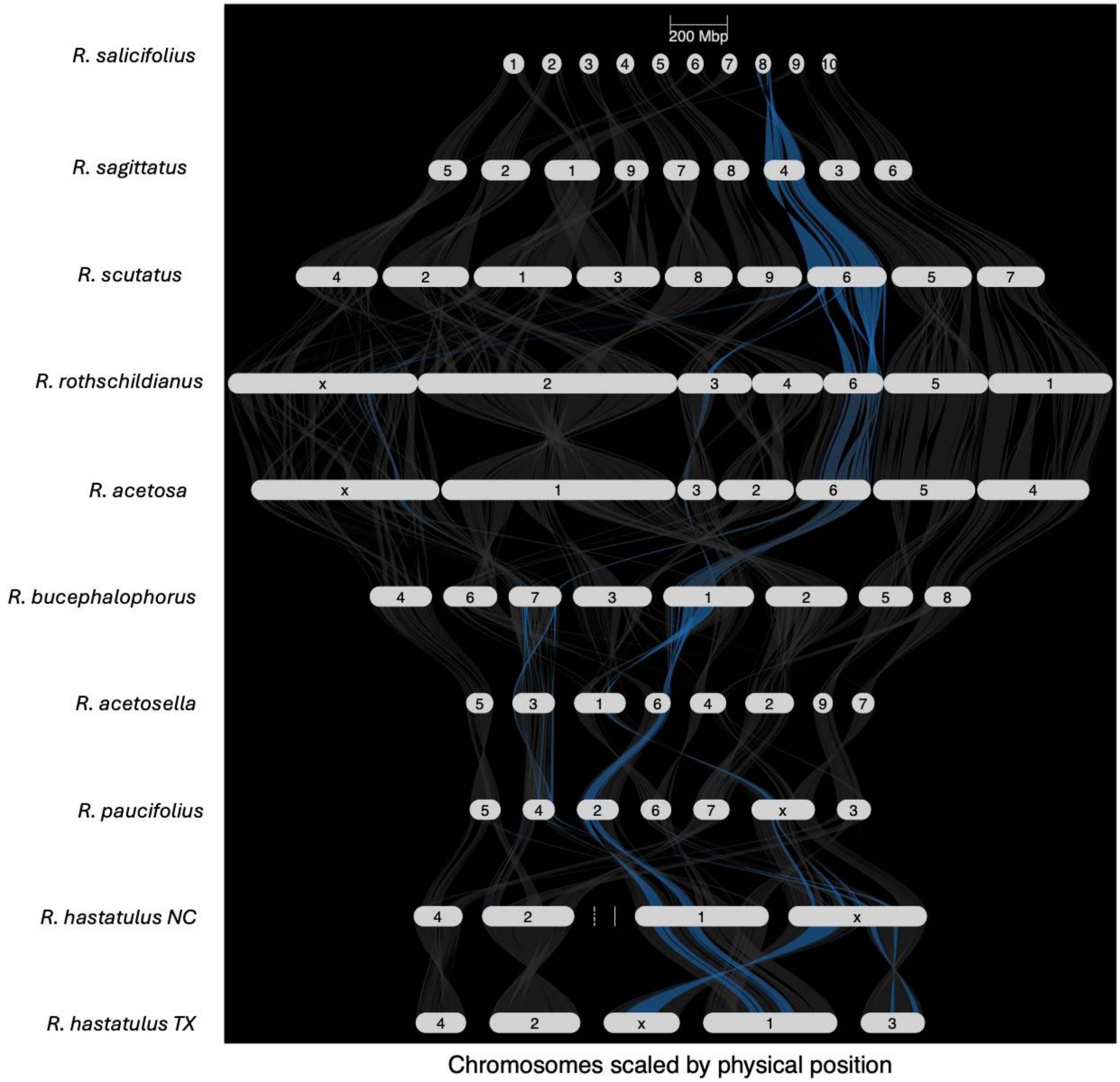


***Supplementary Figure 14.*** Riparian plot depicting synteny between assembled dioecious and hermaphroditic *Rumex* species. The blue bands represent scaffold 8 of the outgroup, *R. salicifolius*.

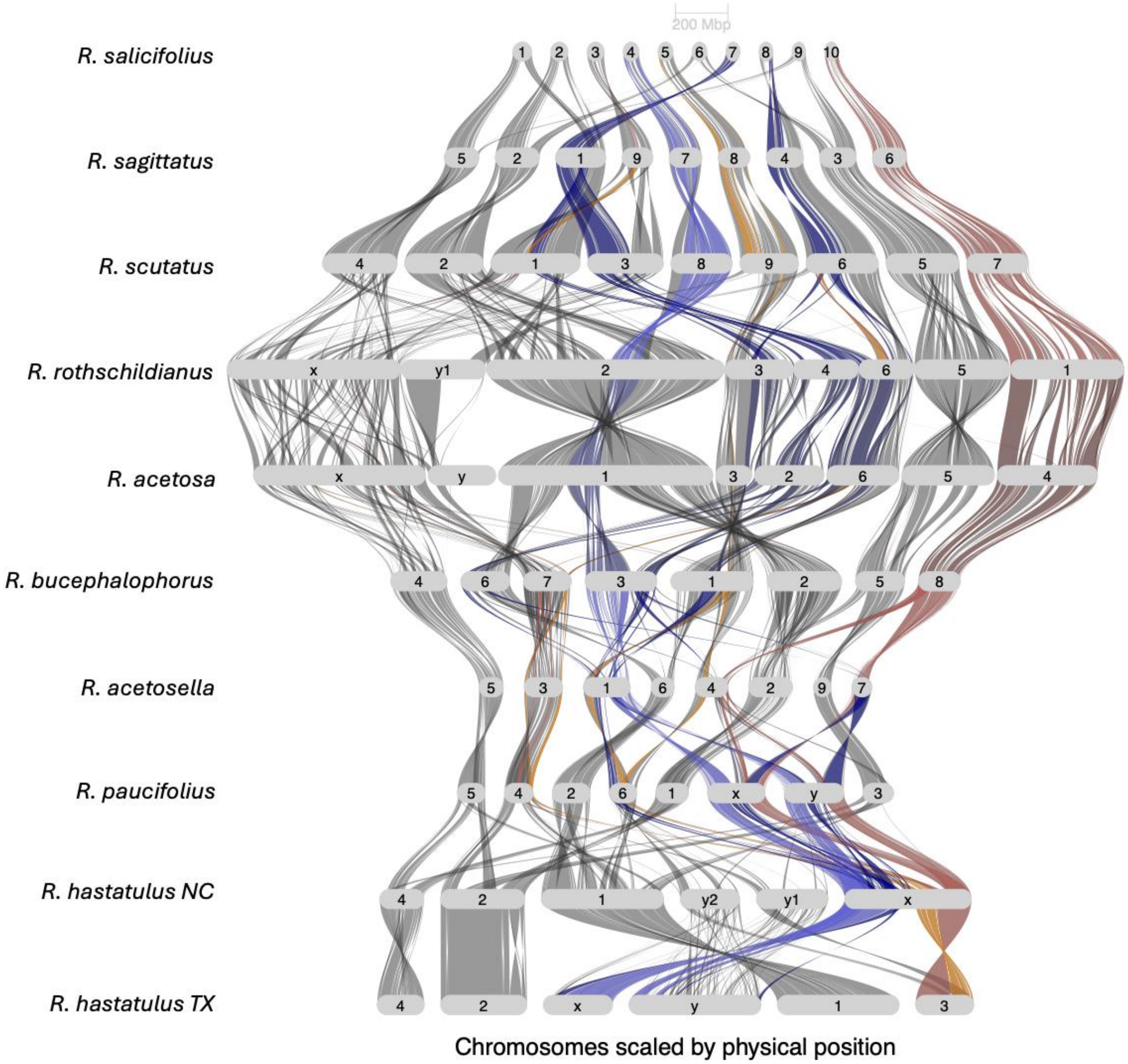


***Supplementary Figure 15.*** Riparian plot depicting synteny between assembled dioecious and hermaphroditic *Rumex* species. The coloured bands depict regions of the X chromosome of the XYY cytotype of *R. hastatulus*. The red bands correspond to the new pseudo-autosomal region, the yellow corresponds to the new sex-linked region, dark blue corresponds to the old-sex linked region, and the light blue corresponds to the old pseudo-autosomal region.

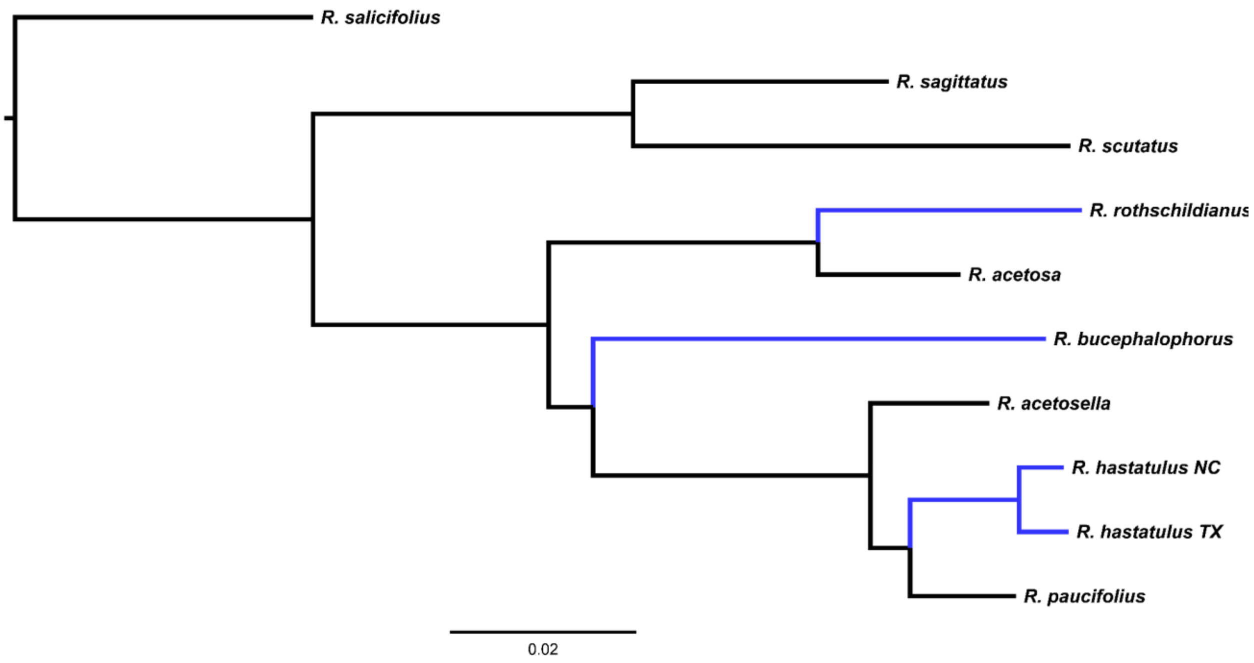


***Supplementary Figure 16***: Phylogeny of *Rumex* inferred from syntenic orthologs with branch lengths in units of substitutions per site. Annual lineages are labelled in blue.

Table 1. Assembly statistics

| Assembly | Assembly length | Contig N50 | Contig L50 | Scaffold N50 | Scaffold L50 | Scaffold N90 | Scaffold L90 | Contig number |
|---|---|---|---|---|---|---|---|---|
| *R. hastatulus* XY maternal | 1,497,422,923 | 7,699,624 | 61 | 315,705,859 | 2 | 173,846,016 | 5 | 1,299 |
| *R. hastatulus* XY paternal | 1,863,807,873 | 5,565,000 | 93 | 465,013,087 | 2 | 179,199,150 | 5 | 4,275 |
| *R. acetosella* haplotype 1 | 2,489,612,908 | 8,895,830 | 82 | 93,788,304 | 10 | 16,278,170 | 27 | 2,578 |
| *R. acetosella* haplotype 2 | 1,418,279,490 | 7,441,330 | 53 | 92,674,719 | 5 | 6,651,151 | 20 | 1,658 |
| *R. acetosa* maternal | 2,874,234,518 | 8,422,662 | 105 | 656,500,594 | 2 | 260,218,433 | 6 | 559 |
| *R. acetosa* paternal | 2,990,395,645 | 7,612,513 | 119 | 353,511,751 | 3 | 15,375,797 | 15 | 555 |
| *R.* | 3,081,491,187 | 3,635,604 | 265 | 662,086,209 | 2 | 248,552,001 | 6 | |

| | | | | | | | | |
|---|---|---|---|---|---|---|---|---|
| *rothschildianus* maternal | | | | | | | | 1,417 |
| *R. rothschildianus* paternal | 2,966,856,971 | 3,699,703 | 238 | 362,463,968 | 3 | 208,556,261 | 7 | 1370 |
| *R. paucifolius* maternal | 908,743,821 | 15,327,942 | 20 | 120,789,685 | 3 | 103,678,334 | 7 | 105 |
| *R. paucifolius* paternal | 932,970,306 | 15,498,446 | 19 | 124,381,532 | 2 | 104,271,472 | 7 | 111 |
| *R. scutatus* | 2,560,304,087 | 53,387,841 | 15 | 278,986,839 | 5 | 224,374,351 | 9 | 2,373 |
| *R. sagittatus* | 1,330,681,304 | 44,852,091 | 11 | 132,305,651 | 5 | 118,223,141 | 9 | 1,071 |

Table 2.

| Assembly | orthoDB | Percent completeness | Complete single copy | Complete duplicates | fragmented | missing |
|---|---|---|---|---|---|---|
| *R. hastatulus* XY maternal | Eukaryota | 97.7 | 110 | 12 | 2 | 1 |
| *R. hastatulus* XY paternal | Eukaryota | 99.2 | 113 | 15 | 1 | 0 |
| *R. hastatulus* XY maternal | Embryophyta | 94.3 | 1809 | 102 | 38 | 77 |
| *R. hastatulus* XY paternal | Embryophyta | 95.1 | 1837 | 90 | 43 | 56 |
| *R. acetosella* haplotype 1 | Eukaryota | 100 | 1 | 128 | 0 | 0 |
| *R. acetosella* haplotype 2 | Eukaryota | 97.7 | 27 | 99 | 0 | 3 |
| *R. acetosella* haplotype 1 | Embryophyta | 97.8 | 15 | 1967 | 13 | 31 |
| *R. acetosella* haplotype 2 | Embryophyta | 94.8 | 413 | 1507 | 29 | 77 |
| *R. acetosa* maternal | Eukaryota | 97.7 | 91 | 35 | 3 | 0 |

| | | | | | | |
|---|---|---|---|---|---|---|
| *R. acetosa* paternal | Eukaryota | 97.7 | 100 | 26 | 0 | 3 |
| *R. acetosa* maternal | Embryophyta | 96.6 | 1703 | 254 | 35 | 34 |
| *R. acetosa* paternal | Embryophyta | 87.1 | 1637 | 128 | 49 | 212 |
| *R. rothschildianus* maternal | Eukaryota | 98.4 | 106 | 21 | 2 | 0 |
| *R. rothschildianus* paternal | Eukaryota | 98.4 | 113 | 14 | 1 | 1 |
| *R. rothschildianus* maternal | Embryophyta | 96.4 | 1796 | 157 | 33 | 40 |
| R. rothschildianus paternal | Embryophyta | 85.5 | 1674 | 59 | 50 | 243 |
| *R. paucifolius* maternal | Eukaryota | 100 | 102 | 27 | 0 | 0 |
| *R. paucifolius* paternal | Eukaryota | 100 | 103 | 26 | 0 | 0 |
| *R. paucifolius* maternal | Embryophyta | 96.3 | 1802 | 149 | 38 | 37 |
| *R. paucifolius* paternal | Embryophyta | 96.2 | 1794 | 155 | 37 | 40 |
| *R. scutatus* | Eukaryota | 99.2 | 113 | 15 | 1 | 0 |
| *R. scutatus* | Embryophyta | 96.7 | 1856 | 104 | 34 | 32 |
| *R. sagittatus* | Eukaryota | 100 | 115 | 14 | 0 | 0 |
| *R. sagittatus* | Embryophyta | 96.2 | 1860 | 89 | 45 | 32 |

Table 3. Results of phylogenetic regression predicting sexual system in *Rumex* species (dioecious vs. hermaphroditic) as a function of the presence/absence of syntenic adjacencies. Adjacency names are based on chromosomal scaffold labels from the assembly of *R. salicifolius*.

| **Variable** | **Metric** | **Value** | **Adjacency** |
|---|---|---|---|
| sexsystemhermaphroditic | p.value | 0.074375 | Scaffold_5Scaffold_8 |

| | | | |
|---|---|---|---|
| sexsystemhermaphroditic | p.value | 0.08021 | Scaffold_8Scaffold_7 |
| sexsystemhermaphroditic | p.value | 0.081804 | Scaffold_3Scaffold_8 |
| sexsystemhermaphroditic | p.value | 0.150976 | Scaffold_3Scaffold_5 |
| sexsystemhermaphroditic | p.value | 0.225281 | Scaffold_2Scaffold_4 |
| sexsystemhermaphroditic | p.value | 0.289338 | Scaffold_8Scaffold_4 |
| sexsystemhermaphroditic | p.value | 0.314921 | Scaffold_3Scaffold_6 |
| sexsystemhermaphroditic | p.value | 0.314921 | Scaffold_5Scaffold_9 |
| sexsystemhermaphroditic | p.value | 0.314921 | Scaffold_8Scaffold_10 |
| sexsystemhermaphroditic | p.value | 0.326359 | Scaffold_1Scaffold_6 |
| sexsystemhermaphroditic | p.value | 0.326359 | Scaffold_2Scaffold_6 |
| sexsystemhermaphroditic | p.value | 0.346988 | Scaffold_1Scaffold_9 |
| sexsystemhermaphroditic | p.value | 0.371563 | Scaffold_7Scaffold_10 |
| sexsystemhermaphroditic | p.value | 0.3724 | Scaffold_7Scaffold_4 |
| sexsystemhermaphroditic | p.value | 0.3824 | Scaffold_3Scaffold_7 |
| sexsystemhermaphroditic | p.value | 0.393999 | Scaffold_1Scaffold_3 |
| sexsystemhermaphroditic | p.value | 0.393999 | Scaffold_1Scaffold_5 |
| sexsystemhermaphroditic | p.value | 0.393999 | Scaffold_2Scaffold_3 |
| sexsystemhermaphroditic | p.value | 0.443006 | Scaffold_1Scaffold_2 |
| sexsystemhermaphroditic | p.value | 0.604945 | Scaffold_2Scaffold_8 |
| sexsystemhermaphroditic | p.value | 0.616006 | Scaffold_1Scaffold_8 |
| sexsystemhermaphroditic | p.value | 0.616006 | Scaffold_5Scaffold_10 |
| sexsystemhermaphroditic | p.value | 0.616006 | Scaffold_4Scaffold_6 |

| | | | |
|---|---|---|---|
| sexsystemhermaphroditic | p.value | 0.624075 | Scaffold_5Scaffold_7 |
| sexsystemhermaphroditic | p.value | 0.624075 | Scaffold_6Scaffold_10 |
| sexsystemhermaphroditic | p.value | 0.631454 | Scaffold_8Scaffold_9 |
| sexsystemhermaphroditic | p.value | 0.817218 | Scaffold_6Scaffold_9 |
| sexsystemhermaphroditic | p.value | 0.860763 | Scaffold_2Scaffold_9 |

Table 4. Genes found both in chromosome 8 of *R. salicifolius* and on one or more X chromosome regions syntenic to chr 8.

| **Gene (*R. salicifolius* *annotation)*** | **X chromosomes** | **Functional annotation** |
|---|---|---|
| NChap2_final_00014329 | *R. rothschildianus* | PRD1 (putative recombination initiation defect 1), *Arabidopsis thaliana* |
| NChap2_final_00015469 | *R. acetosa* | GLCAT14A (beta-glucuronosyltransferase), *Arabidopsis thaliana* |
| NChap2_final_00015504 | *R. acetosa*, *R. rothschildianus* | MFP-a (Glyoxysomal fatty acid beta-oxidation multifunctional protein), *Cucumis sativus* |
| NChap2_final_00018591 | *R. hastatulus* TX | Os09g0364000 (Zinc finger CCCH domain-containing protein), *Oryza sativa* |
| NChap2_final_00018785 | *R. hastatulus* NC | TBL11 (Protein trichome birefringence-like 11), *Arabidopsis thaliana* |
| NChap2_final_00020716 | *R. hastatulus* NC | GYRA (DNA gyrase subunit A%2C chloroplastic/mitochondrial), *Nicotiana benthamiana* |
| NChap2_final_00020729 | *R. acetosella* | SIGB (RNA polymerase sigma factor sigB), |

| | | |
|---|---|---|
| | | *Arabidopsis thaliana* |
| NChap2_final_00021163 | *R. hastatulus* NC | APX6 (Putative L-ascorbate peroxidase 6), *Arabidopsis thaliana* |
| NChap2_final_00021372 | *R. hastatulus* TX | Similar to Glutamine synthetase leaf isozyme chloroplastic, *Phaseolus vulgaris* |
| NChap2_final_00021914 | *R. acetosella*, *R. paucifolius* | NFYA10 (Nuclear transcription factor Y subunit A-10), *Arabidopsis thaliana* |
| NChap2_final_00022877 | *R. hastatulus* TX | CEL3 (endoglucanase 9) *Arabidopsis thaliana* |
| NChap2_final_00022946 | *R. hastatulus* TX | ARF17 (auxin response factor 17), *Arabidopsis thaliana* |
| NChap2_final_00022975 | *R. hastatulus* NC | TMN1 (transmembrane 9 superfamily member 1), *Arabidopsis thaliana* |
| NChap2_final_00022976 | *R. hastatulus* TX | PATL3 (Patellin-3), *Arabidopsis thaliana* |
